# A hyperactivating mutation in formin-like 2 causes brain and vertebrae defects

**DOI:** 10.64898/2026.10.16.759550

**Authors:** Benjamin L. Zaepfel, Joyce Tran, Thomas J. Jucius, Tracy-Ann Read, Grant R. Reeves, Diane Bissen, Eric A. Vitriol, Susan L. Ackerman

**Author notes:** These authors contributed equally to this work (Benjamin L. Zaepfel, Joyce Tran).

## Abstract

Formins are a group of proteins that nucleate and polymerize actin monomers into filaments to promote cell morphology and migration. Mutations in formin-encoding genes cause a broad spectrum of human diseases, ranging from kidney disease to non-syndromic intellectual disability. Through a forward genetics screen, we identified a mouse strain (*nmf418*) with a homozygous mutation in a splice acceptor in *Fmnl2* that causes hindlimb motor dysfunction, enlarged midbrain colliculi, cerebellar hypoplasia, and vertebral abnormalities in the thoracolumbar spinal cord. The *nmf418* mutation causes skipping of exon 10, which encodes a portion of the diaphanous inhibitory domain (DID) required for FMNL2 autoinhibition, and results in dramatically lower FMNL2 protein levels. Consistent with a gain-of-function mechanism, mutant FMNL2 has altered enrichment at filopodia tips, increased actin polymerization activity, and its expression results in enhanced cell migration *in vitro*. Furthermore, none of the neural or skeletal phenotypes were recapitulated in *Fmnl2* knockout mice. Transcriptomic analysis of the embryonic cerebellum suggests that complete loss of *Fmnl2* triggers broad compensatory upregulation of actin regulatory pathways, while expression of mutant FMNL2 is associated with dysregulation of cell migration, motility pathways, and calcium homeostasis pathways. Taken together, these findings suggest that *Fmnl2* is largely dispensable for overall mouse development but that its autoregulation is essential for proper formation of the mouse nervous and skeletal systems.

## Introduction

Mammalian development requires extensive and continuous restructuring of cellular shape to drive cell migration and the formation of organized tissues [1, 2] and specialized cell types [3, 4]. Disruption of this process can impair germ layer specification [5] or body axis formation [6]. The mechanisms that govern changes in cellular structure and cell migration must be tightly regulated to ensure the development of a viable organism.

The cytoskeleton is the central determinant of cellular shape and mobility [7]. It comprises interacting networks of microtubules, intermediate filaments, and actin filaments [8] which provide structural rigidity to withstand physical forces [9] while remaining highly dynamic. Actin filaments have been extensively studied for their roles in cell motility [10–13], muscle contraction [14, 15], and maintenance of cellular shape via the actin cortex [16–18].

Filamentous actin (F-actin) is polymerized from three nearly identical isoforms of globular actin (G-actin) [19]. Linear actin filaments, such as those in filopodia and stress fibers, are primarily nucleated by formins, dimeric proteins defined by a highly conserved formin homology 2 (FH2) domain that is required for nucleation in yeast and is structurally conserved in mice and humans [20–23]. Following nucleation, formins remain associated with the fast-growing barbed end of the filament and drive processive elongation through the direct delivery of profilin-actin complexes by the formin homology 1 (FH1) domain [24, 25].

Diaphanous related formins (DRFs) comprise the largest subfamily of formins containing 10 of the 15 formins in humans [26, 27]. They are regulated by the direct binding of GTP-bound Rho-family GTPases (RhoA, Rac1, Cdc42) to their N-terminal GTPase-binding domain (GBD). This positions DRFs as key effectors of Rho GTPase signaling [28]. In the absence of Rho GTPase binding, access of FH2 to actin monomers is sterically occluded by the intramolecular binding of the diaphanous inhibitory domain (DID), which is adjacent or overlapping with the GBD, to the C-terminal diaphanous autoregulatory domain (DAD) [26, 29, 30]. Inversely, binding of an active Rho GTPase induces conformational changes that relieve DID/DAD autoinhibition and enables DRF dimerization and activity [28, 31, 32].

Mutations in formins have been identified as the monogenic cause of human disease [33–37] and can arise from loss or gain of function. Biallelic variants in *DAAM2* are associated with kidney disease [38]. Homozygous frame-shift mutations in *FMN2* cause nonsyndromic autosomal recessive intellectual disability, and knockout of mouse *Fmn2* reduces dendritic spine density [39]. Homozygous nonsense mutations in the DRF *DIAPH1* have been associated with microcephaly and intellectual disability [39–41]. In contrast, gain-of-function mutations that disrupt the DAD domain in DIAPH1 cause sensorineural hearing loss [42–44]. Activating mutations in the DID domain of diaphanous-related formin family member INF2 cause glomerular degeneration in the kidney, sometimes syndromic with Charcot-Marie-Tooth disease [45, 46]. Together, these findings demonstrate that both decreased and increased formin activity, even from mutations in the same gene, can produce distinct physiological disorders.

Here, we investigate the developmental abnormalities caused by an N-ethyl-N-nitrosourea (ENU)-induced splice mutation in *Fmnl2,* a member of the formin-like (FMNL) subfamily of DRFs. FMNL family members drive profilin-actin-mediated filament elongation but serve as poor actin nucleators [47–49]. This mutation results in a deletion of 25 amino acids from the DID domain, leading to production of a hyperactive FMNL2 variant. Homozygous mutant mice exhibited vertebral defects associated with incomplete spinal column closure and abnormal brain development, whereas *Fmnl2* knockout mice were anatomically normal. *In vitro* experiments revealed that expression of FMNL2 with the DID mutation increased actin polymerization, and fibroblasts from mutant mice had enhanced cellular motility. Together, these results reveal that dysregulated actin dynamics caused by a hyperactive mutation in *Fmnl2* lead to skeletal and brain abnormalities and further highlight that gain-of-function mutations in distinct formins can result in specific and varied pathophysiology.

## Results

### *Nmf418^-/-^* mice exhibit spinal column and brain abnormalities

Mice homozygous for the *nmf418* mutation (*nmf418^-/-^*) were identified by their hindlimb motor defects in an ENU-mutagenesis screen for neurological phenotypes [50]. Gross anatomical analysis of postnatal day 21 (P21) *nmf418^-/-^* mice revealed widening of the spinal column with skin-covered openings reminiscent of closed spina bifida (Fig 1A). *nmf418^-/-^* mice also exhibited expansion of the superior and inferior colliculi and abnormal cerebellar anatomy, including caudal/lateral displacement of the vermis and reduction of the hemispheres (Fig 1B). Histological analysis confirmed cerebellar hypoplasia and abnormal morphology of both the vermis (Fig 1C) and hemispheres (Fig 1D) in *nmf418^-/-^* mice compared to wild-type controls. Additionally, mutant mice displayed severe hypoplasia of the corpus callosum, lateral ventricle enlargement suggestive of hydrocephalus, and ventral displacement of the lateral septal nucleus (Fig 1E).

**Fig. 1.**
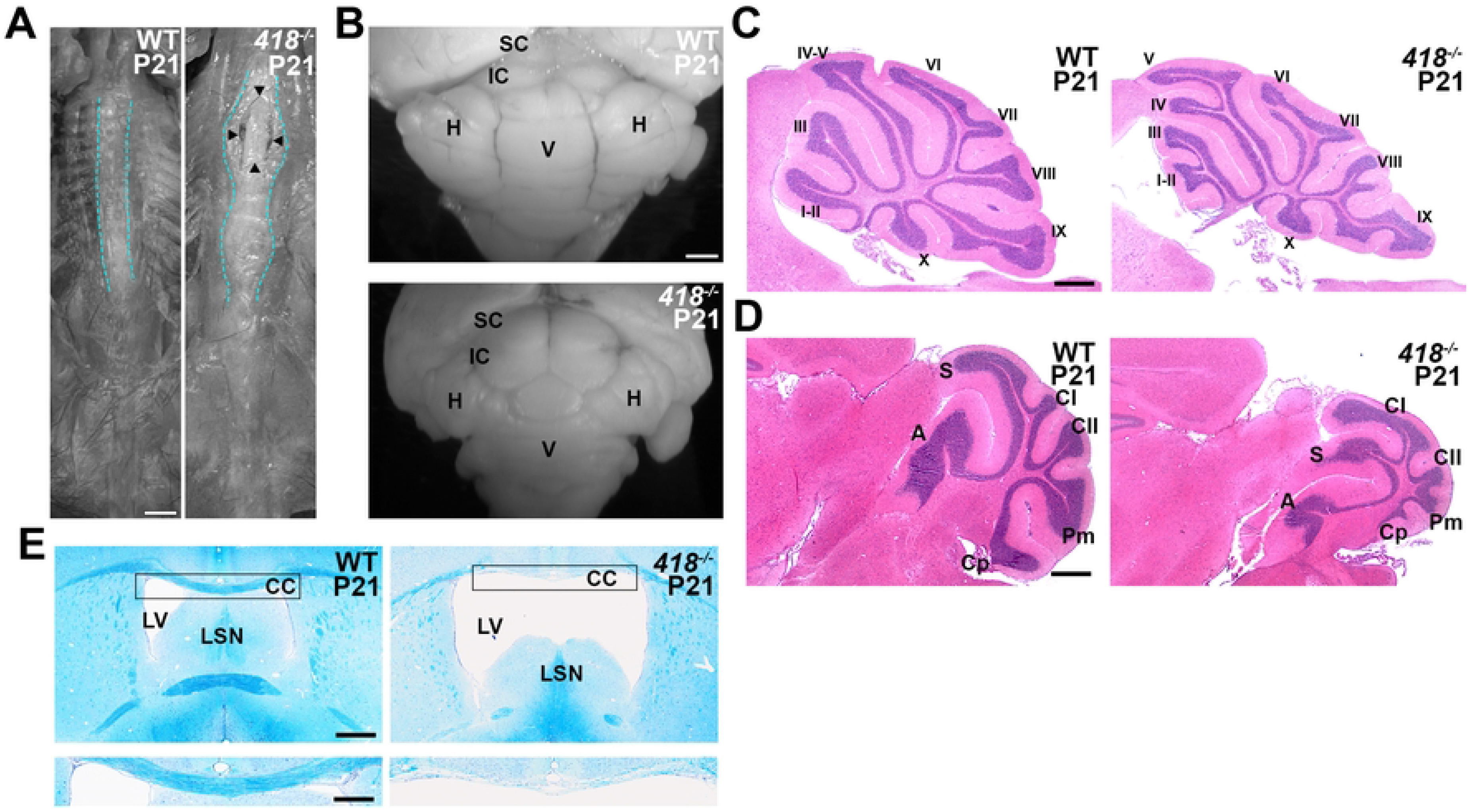
*Nmf418^-/-^* mice have open spinal columns and cerebellar malformations. **A.** Dorsal view of spinal columns from P21 wild-type (WT) and *nmf418*^-/-^ mice. The spinal column boundary is outlined in cyan. Arrowheads mark the boundaries of spinal column opening in *nmf418^-/-^* mice. Scale bar = 5mm. **B.** Caudal view of brains from P21 WT and *nmf418*^-/-^ mice. SC = superior colliculus; IC = inferior colliculus; H = hemisphere; V = vermis. **C, D.** Sagittal sections of cerebellar midline (C) and hemispheres of P21 WT and *nmf418*^-/-^ mice stained with hematoxylin and eosin. Lobules are indicated by Roman numerals (C). Hemispheric lobes (D) are labeled as follows: A, anterior; S, simplex; CI, Crus I; CII, Crus II; Pm, paramedial; Cp, copula pyramidas. Scale bars = 200μm. **E.** Coronal sections from P21 WT and *nmf418*^-/-^ brains stained with cresyl violet/luxol fast blue. Note the enlarged lateral ventricles (LV), deviated lateral septal nucleus (LSN, upper panel) and hypoplastic corpus callosum (CC, lower panel) in mutant mice. Scale bars = 500μm (upper) and 200μm (lower).

### *Nmf418* is a splicing mutation in *Formin-like 2*

The *nmf418* mutation was induced on a C57BL/6J background and initially mapped to Chromosome 2 in a cross to BALB/cByJ mice. Fine-mapping by recombination analysis of F2 mice from a C57BL/6J; BALB-*nmf418^+/-^* x NOD/ShiLtJ cross localized the mutation to a 4-Mb interval between *D2Mit458* and *rs27894839*, a region containing 16 protein-coding genes (Fig 2A).

**Fig. 2.**
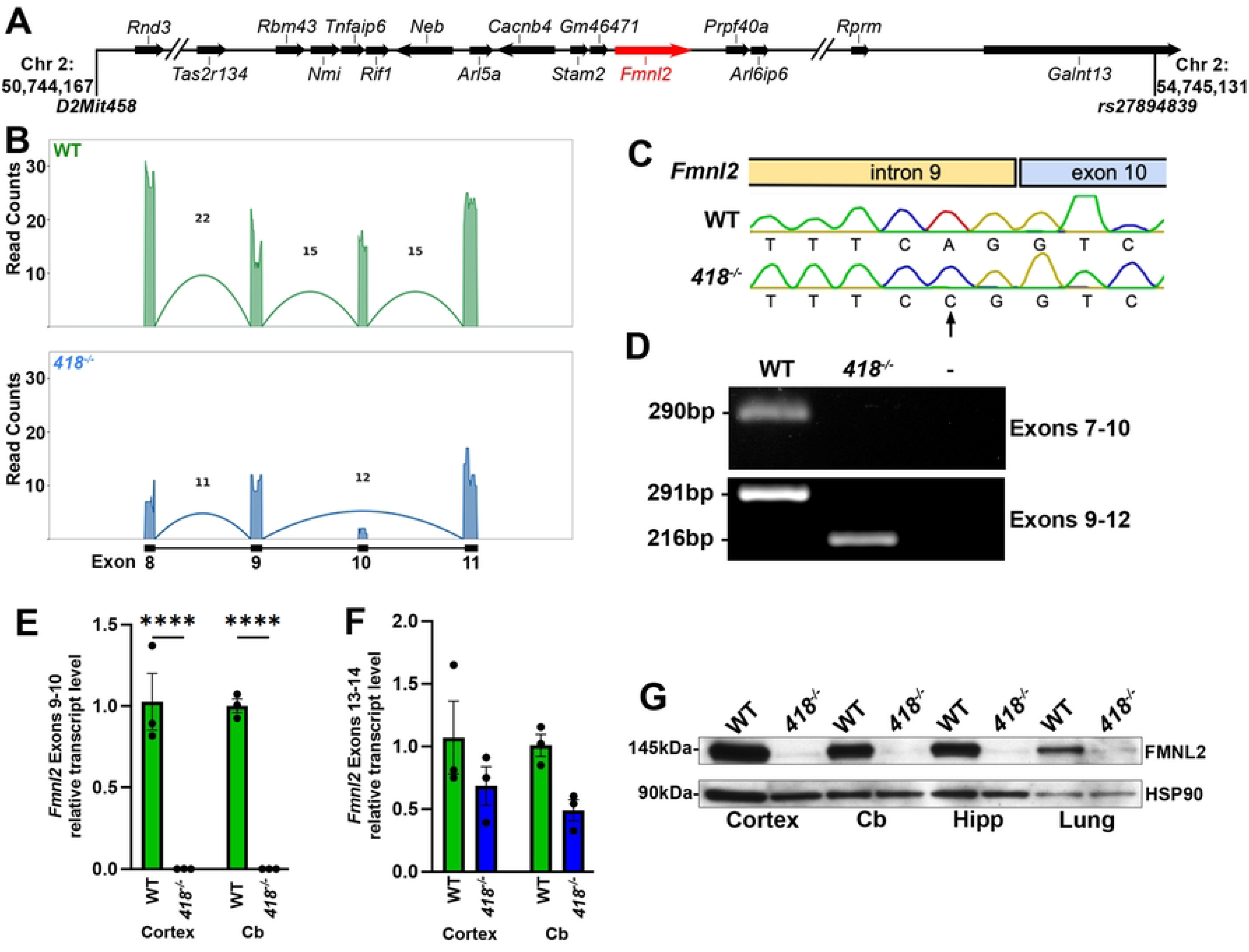
The *nmf418* mutation causes in-frame skipping of *Fmnl2* exon 10. **A.** Schematic of the *nmf418* critical region on Chromosome 2 bounded by markers *D2Mit458* and *rs27894839*. Reference genome is GRCm39. **B.** Representative sashimi plots of *Fmnl2* exons 8-11 from RNA-Seq data from wild-type and *nmf418^-/-^* E18.5 cerebella. Numbers above arcs indicate exon-exon junction reads for the representative library. Note the lack of exon 10 junction reads and the presence of these reads for exon 9-11 in mutant. **C.** Chromatograms of *Fmnl2* intron 9/exon 10 boundaries in wild-type and *nmf418^-/-^* samples. Arrow denotes the A>C transversion at the canonical AG splice acceptor in intron 9. **D.** RT-PCR amplicons spanning *Fmnl2* exons 7-10 (upper) and 9-12 (lower). Gels were stained with ethidium bromide. - = no template control. **E, F.** Quantitative RT-PCR analysis of *Fmnl2* transcripts containing exon 9-10 junctions (**E**) or exon 13-14 junctions (**F**). Data were normalized to *Gapdh* and transcript levels are relative to that of wild type mice. Data are means ± SEM. **G.** Representative western blot of mouse tissue extracts for FMNL2. HSP90 was used as loading control. **** = p < 0.0001; two-way ANOVA with Šidák’s multiple comparisons test. Cb = cerebellum, Hipp = hippocampus.

To identify the causal variant, we isolated RNA from the cerebella of E18.5 embryos that were presumed (and later confirmed) to be wild-type or *nmf418^-/-^* based on the absence or presence of spinal column openings and performed RNA sequencing. Transcripts from 13 of the 16 genes in the critical region were detected; neither expression differences nor coding sequence polymorphisms were identified. However, exon 10 of *Fmnl2* (Formin-like 2), which is retained in wild-type transcripts, was skipped in mutant transcripts (Fig 2B). Sanger sequencing of genomic DNA at the border of intron 9 and exon 10 revealed that the canonical AG splice acceptor site of intron 9 was altered to CG in the mutant genome (Fig 2C). We confirmed exon 10 skipping by sequencing RT-PCR amplicons generated from wild-type and mutant cerebellar cDNA using primers within exons 7/10 and exons 9/12 (Fig 2D). Consistently, qRT-PCR detected exon 10-containing *Fmnl2* transcripts in wild-type but not mutant cortex and cerebellum (Fig 2E).

Exon 10 is 75 nucleotides; hence its skipping is predicted to preserve the reading frame and not trigger nonsense-mediated decay. Consistent with this, while trending downward, overall *Fmnl2* transcript levels were not statistically different between genotypes by qRT-PCR using exon 13-14 primers (Fig 2F). However, a marked reduction in FMNL2 protein in mutant tissues was observed by western blot analysis (Fig 2G).

To confirm causality, we used CRISPR-Cas9 to introduce the intron 9 A>C mutation into wild-type mice (*Fmnl2^In9A>C/In9A>C^*; Fig S1A). These mice recapitulated the hindlimb motor defects, cerebellar hypoplasia and disorganization (Fig S1B), and the dramatic reduction in FMNL2 protein levels (Fig S1C) observed in *nmf418*^-/-^ mice. Together, these data confirm that the A>C mutation at the *Fmnl2* intron 9/exon 10 splice junction is the causative mutation in *nmf418*^-/-^ mice, hereafter referred to textually as *Fmnl2^418/418^*.

### FMNL2Δ25 has increased activity

FMNL2 function, like that of other diaphanous-related formins, is repressed by intramolecular autoinhibition until an active Rho GTPase binds the GBD [30]. Autoinhibition is mediated by intramolecular binding of the C-terminal DAD to the N-terminal DID, which includes the GBD and formin homology domain 3 (FH3). The FH3 domain of FMNL2 comprises five armadillo repeat motifs (ARMs). Exon 10 of *Fmnl2* encodes a portion of the fourth ARM which has been shown to interact closely with CDC42 [51] to promote release of autoinhibition (Fig 3A, B). We therefore hypothesized that loss of this exon in *Fmnl2^418/418^* mice either (1) disrupts FMNL2-CDC42 complex formation and prevents release of DID/DAD autoinhibition or (2) prevents DID/DAD interaction independent of CDC42 binding. The former could lead to hypofunctional protein due to extended or constitutive autoinhibition, while the latter could lead to hyperactivity by reducing autoinhibitory DID/DAD interactions.

**Fig. 3.**
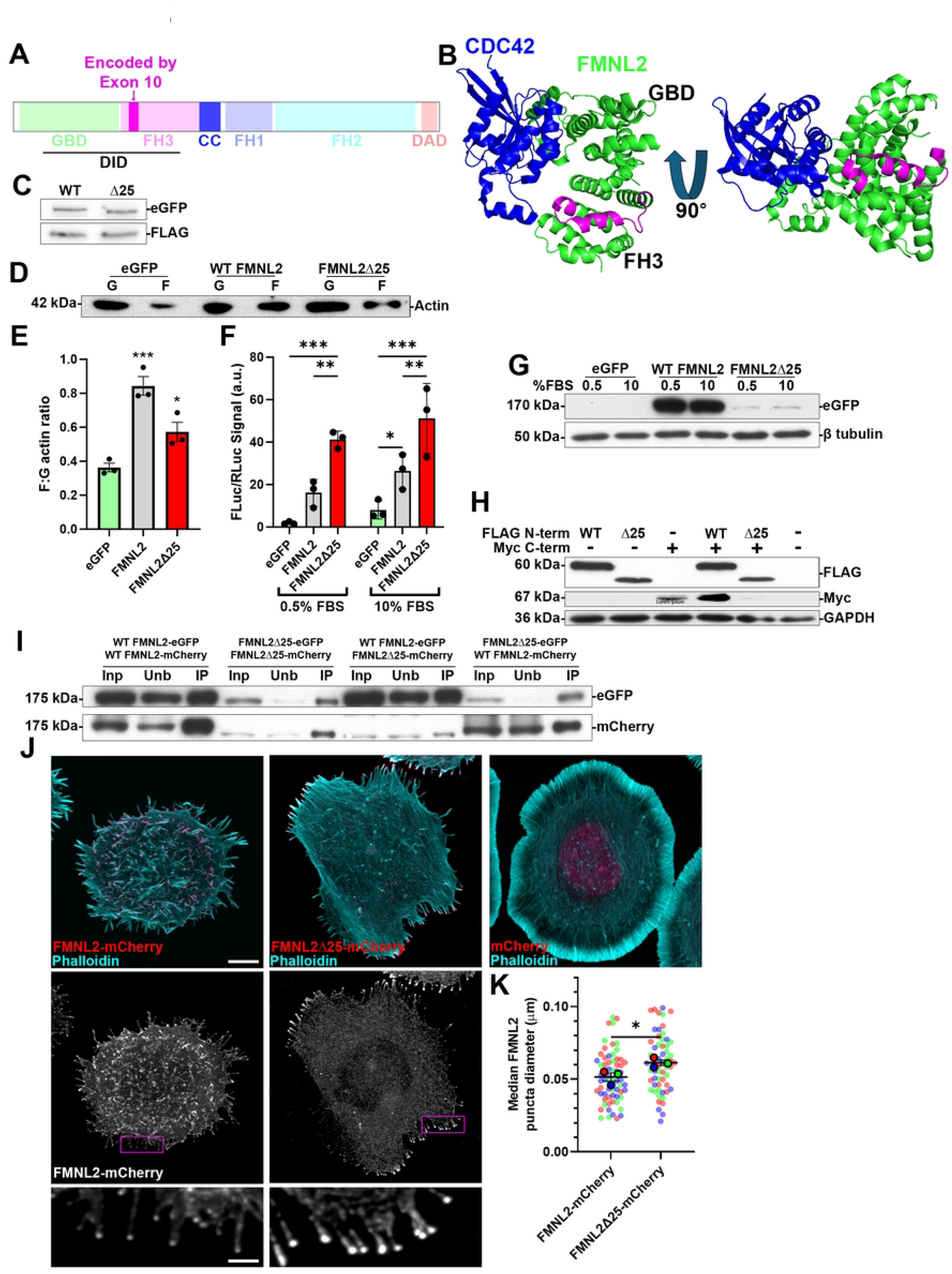
Loss of exon 10 results in hyperactive FMNL2. **A.** Schematic illustrating FMNL2 protein domains. GBD, GTPase-binding domain; FH3, formin homology 3 domain; CC, coiled coil domain; FH1, formin homology 1 domain; FH2, formin homology 2 domain; DAD, Diaphanous autoregulatory domain; DID, Diaphanous inhibitory domain. The region encoded by exon 10 within the FH3 domain is shown in magenta. **B.** Representation of the crystal structure of the FMNL2 N-terminus (green) in complex with CDC42 (blue) [51]. Residues encoded by *Fmnl2* exon 10 are shown in magenta. Note the proximity of CDC42 to amino acids encoded by exon 10. **C.** Representative western blot of immunoprecipitated wild-type FMNL2-eGFP (WT) and FMNL2Δ25-eGFP (Δ25) and co-precipitated FLAG-CDC42 G12V. **D.** Representative western blot of globular (G) and filamentous (F) actin from fractionated HEK293T cells overexpressing eGFP, wild-type (WT) FMNL2-eGFP, or FMNL2Δ25-eGFP. The blot was probed with actin antibodies. **E.** Quantification of F:G actin ratios from three independent fractionation experiments. Statistics calculated by one-way ANOVA with Dunnett’s multiple comparisons test (all conditions compared to eGFP). Data are means ± SEM. **F.** Firefly (Fluc):*Renilla* (Rluc) luciferase ratio from HEK293T cells co-transfected with SRE-Fluc and Rluc reporters and plasmids encoding eGFP, FMNL2-eGFP, or FMNL2Δ25-eGFP at 0.5% and 10% fetal bovine serum (FBS). Statistics calculated by two-way ANOVA with Tukey’s multiple comparisons test. Data are means ± SEM. **G.** Representative western blot of eGFP and β-tubulin loading control from the corresponding samples in (**F**) grown in 0.5% or 10% serum. **H.** Representative western blot of samples from HEK293T cells for eGFP and overexpressing the noted FMNL2 N-terminal (3xFlag) and C-terminal (3xMyc) variants tagged at the N-terminus. Blots were probed for FLAG- and Myc-epitope tags and GAPDH as loading control. Note that the levels of the C-terminal fragment are increased upon expression of the wild type N-terminal fragment but decreased when co-expressed with the mutant fragment. **I.** Representative co-immunoprecipitation western blot showing input (Inp), unbound (Unb) fraction, and eGFP-immunoprecipitated (IP) fraction from HEK293T cells overexpressing the indicated eGFP- and mCherry-tagged FMNL2 variants. Blots were probed for eGFP (bait) and mCherry (prey). Note that each FMNL2 variant co-precipitates itself and the other variant. **J.** Representative images of CAD cells overexpressing FMNL2-mCherry, FMNL2Δ25-mCherry, or mCherry alone (control). Top row: merged images of mCherry (magenta) and phalloidin-stained F-actin (cyan). Scale bar = 5μm. Middle row: mCherry channel alone. Bottom row: magnified insets of boxed regions on middle row showing filopodia tip localization. Scale bar = 1μm. **K.** Quantification of the median FMNL2-mCherry puncta diameter at filopodia tips. Small dots represent individual cell medians, outlined dots represent replicate means. Replicates are color coded. Note that FMNL2Δ25-mCherry puncta are significantly larger, consistent with increased tip enrichment. Statistics were calculated by paired t-test. * = p < 0.05; ** = p < 0.01; *** = p < 0.001.

To test whether loss of the 25 amino acids encoded by exon 10 in FMNL2 (FMNL2Δ25) alters CDC42 interactions, we co-immunoprecipitated eGFP-tagged wild-type FMNL2 or FMNL2Δ25 following co-expression with constitutively active Flag-tagged CDC42 (CDC42 G12V) in HEK293T cells. Coprecipitation of FLAG-CDC42 G12V was comparable with wild-type FMNL2 and FMNL2Δ25 (Fig 3C), suggesting that although ARM four of FMNL2 structurally interacts with CDC42 [51], this interaction may not contribute to the overall binding affinity between these two proteins.

We next examined the impact of exon 10 loss on actin polymerization activity by measuring the F-actin:G-actin ratio in HEK293T cells overexpressing wild-type FMNL2-eGFP, FMNL2Δ25-eGFP, or eGFP alone (Figs 3D, E). Both FMNL2-eGFP and FMNL2Δ25-eGFP increased the F-actin:G-actin ratio relative to eGFP controls, indicating that FMNL2Δ25 retains actin polymerization activity. Consistent with observations in *Fmnl2^418/418^* tissues, FMNL2Δ25-eGFP was expressed at lower levels than wild-type FMNL2-eGFP (Fig S2A). When the F-actin:G-actin ratio was normalized to eGFP-tagged protein levels, expression of FMNL2Δ25-eGFP produced an approximately five-fold greater ratio than wild-type FMNL2-eGFP (Fig S2B), indicating that FMNL2Δ25 has substantially higher actin polymerization activity per molecule.

We corroborated these findings using a serum response element (SRE) luciferase reporter assay, which measures actin polymerization indirectly through the MAL/SRF transcriptional pathway [52]. In this assay, actin polymerization reduces the free G-actin pool, releasing MAL from cytoplasmic sequestration by G-actin and enabling its nuclear translocation and activation of SRE-driven transcription (Fig S3). HEK293T cells were co-transfected with an SRE-firefly luciferase reporter, a *Renilla* luciferase normalization construct, and plasmids encoding wild-type FMNL2-eGFP or FMNL2Δ25-eGFP. Both constructs increased the firefly:*Renilla* luciferase ratio relative to eGFP controls in a serum-independent manner (Fig 3F). Notably, FMNL2Δ25-eGFP produced a greater increase than wild-type FMNL2-eGFP despite significantly lower protein expression (Fig 3F, G). Together, these results demonstrate that loss of exon 10 confers hyperactive actin polymerization activity by FMNL2.

A recent study of the formin DIAPH3 suggests that the DID/DAD interaction promotes protein stability [53]. To interrogate the possibility that a decrease in this interaction in FMNL2Δ25 underlies its reduced abundance, we co-transfected HEK293T cells with wild-type N-terminus or Δ25 N-terminal fragments, (both N-terminally tagged with 3xFlag) and the shared C-terminal fragment (N-terminally tagged with 3xMyc), alone or in combination. Expression of wild-type N-terminus increased C-terminal fragment abundance, consistent with DIAPH3 findings (Fig 3H). In contrast, co-expression of Δ25 N-terminus reduced C-terminal abundance, suggesting that loss of the residues encoded by exon 10 destabilizes the DID/DAD interaction and thereby reduced FMNL2 protein stability in *Fmnl2^418/418^* mouse tissues.

Since diaphanous-related formins dimerize via a parallel coiled-coil motif C-terminal to the FH3 domain to promote processive actin assembly, we asked whether FMNL2Δ25 retains dimerization capacity. We co-transfected HEK293T cells with wild-type FMNL2 or FMNL2Δ25, each tagged at the C-terminus with eGFP or mCherry and assessed interactions by co-immunoprecipitation and western blot. Wild-type FMNL2-eGFP pulled down wild-type FMNL2-mCherry, and co-immunoprecipitation was similarly observed between FMNL2Δ25 variants and between wild-type FMNL2 and FMNL2Δ25 (Fig 3I). These results demonstrate that loss of the 25 amino acids encoded by exon 10 does not disrupt FMNL2 dimerization.

We next examined the effect of FMNL2Δ25 on filopodia, actin-based membrane protrusions requiring the coordinated assembly of long, unbranched actin filaments by a formin-containing complex at their tips [54]. Expression of wild-type FMNL2-mCherry in Cath.a-differentiated (CAD) cells [55] increased filopodia formation distributed uniformly around the cell perimeter compared to mCherry controls, consistent with previous reports [48, 56]. In contrast, FMNL2Δ25-mCherry induced filopodia in a more polarized distribution at opposing cellular edges, and cells expressing FMNL2Δ25-mCherry also displayed a flattened, reticular morphology (Fig 3J). Consistent with constitutively active FMNL2 [48, 57–60], FMNL2Δ25-mCherry showed enriched localization at filopodia tips with markedly reduced cytoplasmic signal, whereas wild-type FMNL2-mCherry was distributed throughout the cell body (Fig 3J, K). Collectively, our observations of increased F-actin:G-actin ratio, elevated SRE reporter activity, destabilization of FMNL2Δ25, and filopodia tip-enriched localization support the conclusion that loss of exon 10 renders FMNL2 hyperactive.

### Loss of function of *Fmnl2* does not recapitulate *Fmnl2^418/418^* phenotypes

The data above suggest that the *nmf418* mutation renders FMNL2 hyperactive. However, since FMNL2 levels are reduced in *Fmnl2^418/418^* tissues, the observed phenotypes could also reflect loss of FMNL2 function. To distinguish between these possibilities, we generated a conditional knockout mouse with loxP sites flanking exon 4 of *Fmnl2* (Figure S4A). Deletion of this 77-nt exon causes a frameshift and premature stop codon, resulting in complete absence of FMNL2 protein in mice expressing Sox2-Cre (*Fmnl2^-/-^*; Fig S4B).

*Fmnl2*^-/-^ mice were recovered at P21 at normal Mendelian ratios, whereas only ∼4% of *Fmnl2^418/418^* and *Fmnl2*^In9A>C/In9A>C^ mice survived to P21, despite being present at Mendelian ratios at E18.5 (Fig S4C). This early lethality in hyperactive, but not null, mice further support that the *Fmnl2^418^* is not a loss-of-function mutation.

To further test this, we examined *Fmnl2*^-/-^ mice for additional phenotypes that were present in *Fmnl2^418/418^* mice. *Fmnl2*^-/-^ mice showed no gross motor defects at P21 and no openings in the spinal column at E18.5 (Fig 4A). In contrast, spinal column openings extending from the midthoracic spinal cord to the first sacral segment were present in all *Fmnl2^418/418^* (14/14) and *Fmnl2^In9A>C/In9A>C^* (11/11) embryos examined at this age (Fig 4A). Closed forms of spina bifida are frequently caused by one or more improperly formed vertebrae [61], whereas more severe forms of this disorder are often a result of failure of the neural tube to completely close, which normally occurs by E10 in mice [62]. Indeed, when we examined horizontal sections along the anterior-posterior axis of E11.5 embryos, the neural tube was closed in *Fmnl2^418/418^* embryos (Fig S5A). However, Alcian blue/Alizarin red skeletal staining of E18.5 embryos revealed widening of the vertebral column along the medio-lateral axis at thoracic and lumbar levels in *Fmnl2^418/418^* mutants, with the most pronounced defects in spinous process fusion at L1 (Fig 4B). E18.5 *Fmnl2^418/418^* lumbar spinal cords appeared flattened compared to wild-type, corresponding to the region with the greatest spinal column widening (Fig S5B). No vertebral abnormalities were observed in *Fmnl2*^-/-^ or wild-type mice (Fig 4A, B).

**Fig. 4.**
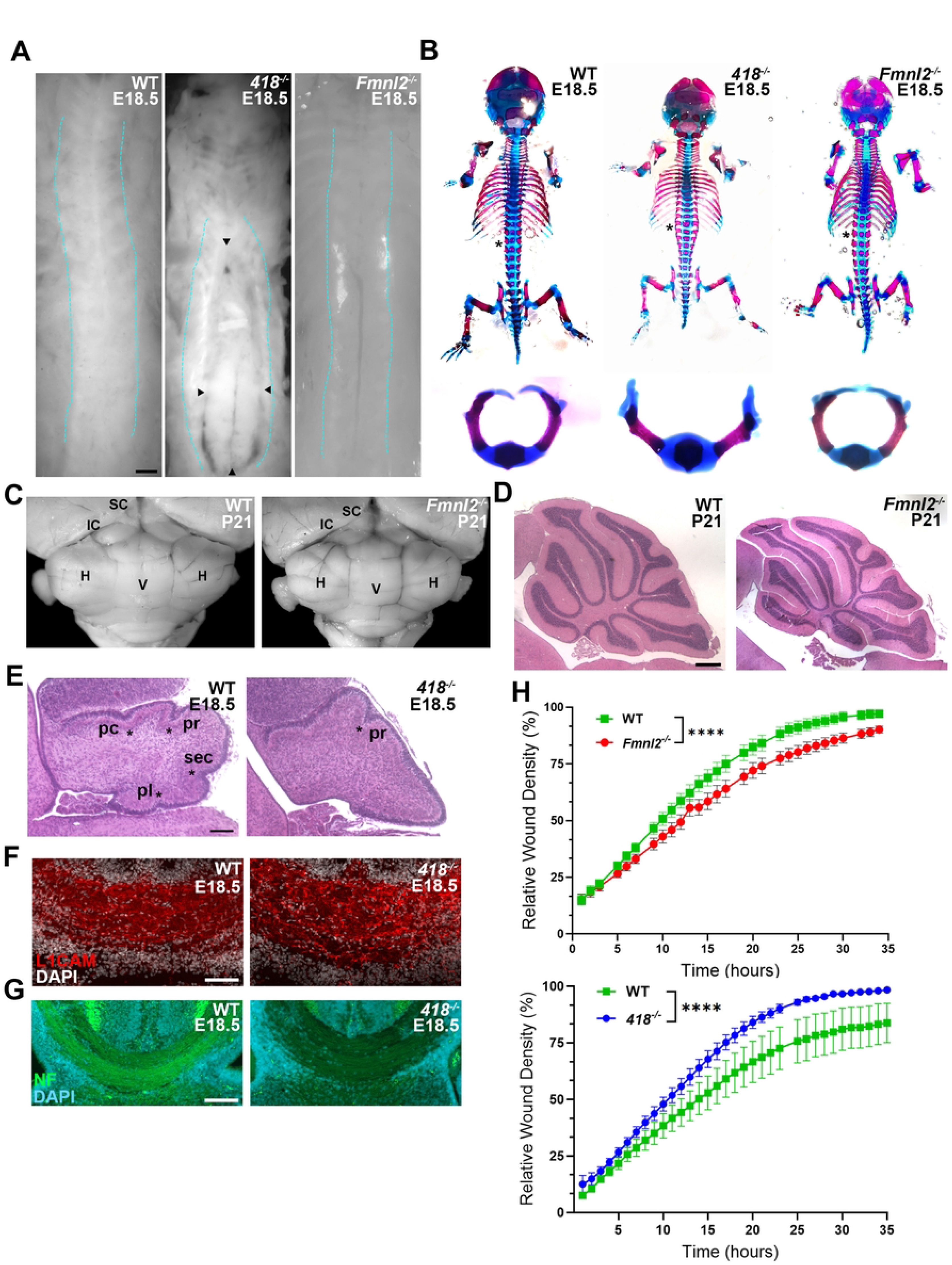
*Fmnl2^418^* induces spinal, cerebellar, and cell motility changes distinct from *Fmnl2* loss-of-function. **A.** Dorsal view of spinal columns from E18.5 wild-type (WT), *Fmnl2^418/418^* (*418*^-/-^), and *Fmnl2*^-/-^ mice. Cyan lines delineate spinal column boundaries. Black arrowheads denote boundaries of spinal column opening in *Fmnl2^418/418^* mice. **B.** Dorsal views of alcian blue (cartilage)/alizarin red (bone)-stained skeletons from WT, *Fmnl2^418/418^,* and *Fmnl2*^-/-^ E18.5 embryos (top panels). Asterisks denote L1 vertebrae shown at higher magnification (lower panels). Note the widening of the vertebral column in *Fmnl2^418/418^* mice and the lack of closure of the vertebral dorsal spinous process. **C.** Caudal views of cerebella from P21 WT and *Fmnl2*^-/-^ mice. SC = superior colliculus; IC = inferior colliculus; H = hemisphere; V = vermis. **D.** Midline sagittal sections of P21 cerebella from WT and *Fmnl2*^-/-^ mice stained with hematoxylin and eosin. Scale bar = 200μm. **E.** Midline sagittal sections of cerebella from E18.5 WT and *Fmnl2^418/418^* mice stained with hematoxylin and eosin. Asterisks denote visible principal fissures, several of which are not present in *Fmnl2^418/418^* at this age. pc, preculminate fissure; pr, primary fissure; sec, secondary fissure; pl, posterolateral fissure. Scale bar = 100μm. **F, G.** Immunofluorescence using antibodies to L1CAM (F) and pan-phospho-neurofilament (NF, panel G) of the corpus callosum in coronal sections of E18.5 WT and *Fmnl2^418/418^* mice. Sections are counterstained with DAPI. Scale bars = 100μm (F) and 200μm (G). Note the corpus callosum is not hypoplastic at E18.5. **H.** Relative wound density over 35 hours for WT and *Fmnl2^-/-^* and WT and *Fmnl2^418/418^* mouse embryonic fibroblasts (MEFs) following scratch wounding. Data are means ± SEM from three independent experiments. *Fmnl2*^-/-^ MEFS show slower gap closure than WT whereas *Fmnl2^418/418^* MEFs have faster gap closure. Statistics calculated by two-way ANOVA. **** indicates p < 0.0001.

Cerebellar defects were similarly absent in P21 *Fmnl2*^-/-^ mice, whereas *Fmnl2^418/418^* embryos displayed flattening of the dorsal cerebellum and enlargement of the inferior colliculus by E18.5 (Fig 4C-E). Given that formins have been shown to govern axon growth cone behavior, we examined corpus callosal axons at midline in E18.5 *Fmnl2^418/418^* and wild-type brains by immunofluorescence using antibodies against L1CAM (Fig 4F) or a pan-phospho-neurofilament antibody (Fig 4G) [63]. Unlike at P21, the width of this commissure was similar between wild-type and *Fmnl2^418/418^* embryos, indicating that midline crossing is not disrupted in *Fmnl2^418/418^* mice (Fig 4 F, G). This finding also suggests that the corpus callosum hypoplasia observed postnatally arises from degeneration of these axons (Figs 1E). Interestingly, immunofluorescence intensity was reduced in mutant brains, suggesting that corpus callosal axons in *Fmnl2^418/418^* mice may undergo delayed maturation [64, 65].

We examined whether mice carrying a single hyperactive allele would exhibit any obvious neural phenotypes as would be expected of a dominant gain-of-function variant. Unlike the postnatal lethality of *Fmnl2^418/418^* mice, the expected ratio of *Fmnl2^+/418^* mice was observed at P21 (Fig S4C). At E18.5, *Fmnl2^+/418^* mice exhibited no obvious skeletal, cerebellar, or spinal defects (Fig S5C-E), nor did we observe any gross abnormalities in these structures at P21 (Fig S5F). These findings suggested that the presence of the more highly expressed wild-type FMNL2 is sufficient to regulate the mixed pool of FMNL2. Furthermore, we did not observe cerebellar hypoplasia (Fig S5G) or early lethality (Fig S5H) in P21 mice carrying a single mutant and knockout allele (*Fmnl2^418/-^*). Together, these findings suggest that although FMNL2Δ25 exhibits hyperactivity, the low expression from a single allele is not sufficient to elicit the mutant phenotypes.

To determine whether FMNL2Δ25 expression or loss of FMNL2 affects cell motility, we performed wound healing assays using wild-type, *Fmnl2^-/-^*, and *Fmnl2^418/418^* mouse embryonic fibroblasts (MEFs). As expected, FMNL2 abundance was markedly reduced in *Fmnl2^418/418^* MEFs relative to wild type (Fig S6A). Consistent with previous reports that loss of diaphanous formins reduces motility in adherent fibroblasts [66], *Fmnl2^-/-^* MEFs showed slower gap closure than wild-type MEFs (Figs 4H, S6B). In contrast, gap closure was faster in *Fmnl2^418/418^* MEFs than wild-type MEFs (Figs 4H, S6C). To confirm that these differences reflected motility rather than proliferation, we assessed cell cycle status by BrdU incorporation and Ki67 immunocytochemistry. No differences in the proportion of cells in active phases of the cell cycle (Ki67+) or S-phase (BrdU+) were detected across genotypes (Fig S6D).

### Hyperactive FMNL2 drives distinct transcriptional changes from FMNL2 loss

Finally, to examine potential transcriptional distinctions between *Fmnl2^418/418^* and *Fmnl2^-/-^* mice, we analyzed RNA-Seq data from the cerebellum of E18.5 wild-type, *Fmnl2^418/418^*, and *Fmnl2^-/-^* embryos (Fig S7A). Loss of *Fmnl2* resulted in more differentially expressed genes (DEGs; 2086 upregulated and 2253 downregulated relative to WT) than the *Fmnl2^418^* mutation (247 upregulated and 551 downregulated) with some overlap between genotypes (Fig S7B). Principal component analysis further suggested that the two alleles differentially affect gene regulation, with each genotype clustering separately (Fig S7C).

Gene ontology (GO) analysis of DEGs between wild-type and *Fmnl2^418/418^* cerebella revealed downregulation of genes associated with “small GTPase mediated signal transduction”, consistent with dysregulated signaling downstream of the FMNL2-CDC42 interaction (Fig 5A, Table S1). The “cell migration” term was enriched among downregulated genes in both *Fmnl2^418/418^* and *Fmnl2^-/-^* data sets. Interestingly, transcripts involved in regulation of ryanodine-sensitive calcium release were upregulated in the *Fmnl2^418/418^* cerebellum. Ryanodine receptors occupy microdomains on the endoplasmic reticulum (ER) membrane and are directly tethered to voltage-gated calcium channels on the plasma membrane [67], and the actin cytoskeleton regulates the anterograde trafficking, membrane tethering, clustering, and microdomain anchoring of these proteins [68, 69]. Hyperactive actin polymerization driven by FMNL2Δ25 could therefore disrupt the spatial positioning of these channels, potentially altering calcium signaling in cerebellar neurons.

**Fig. 5:**
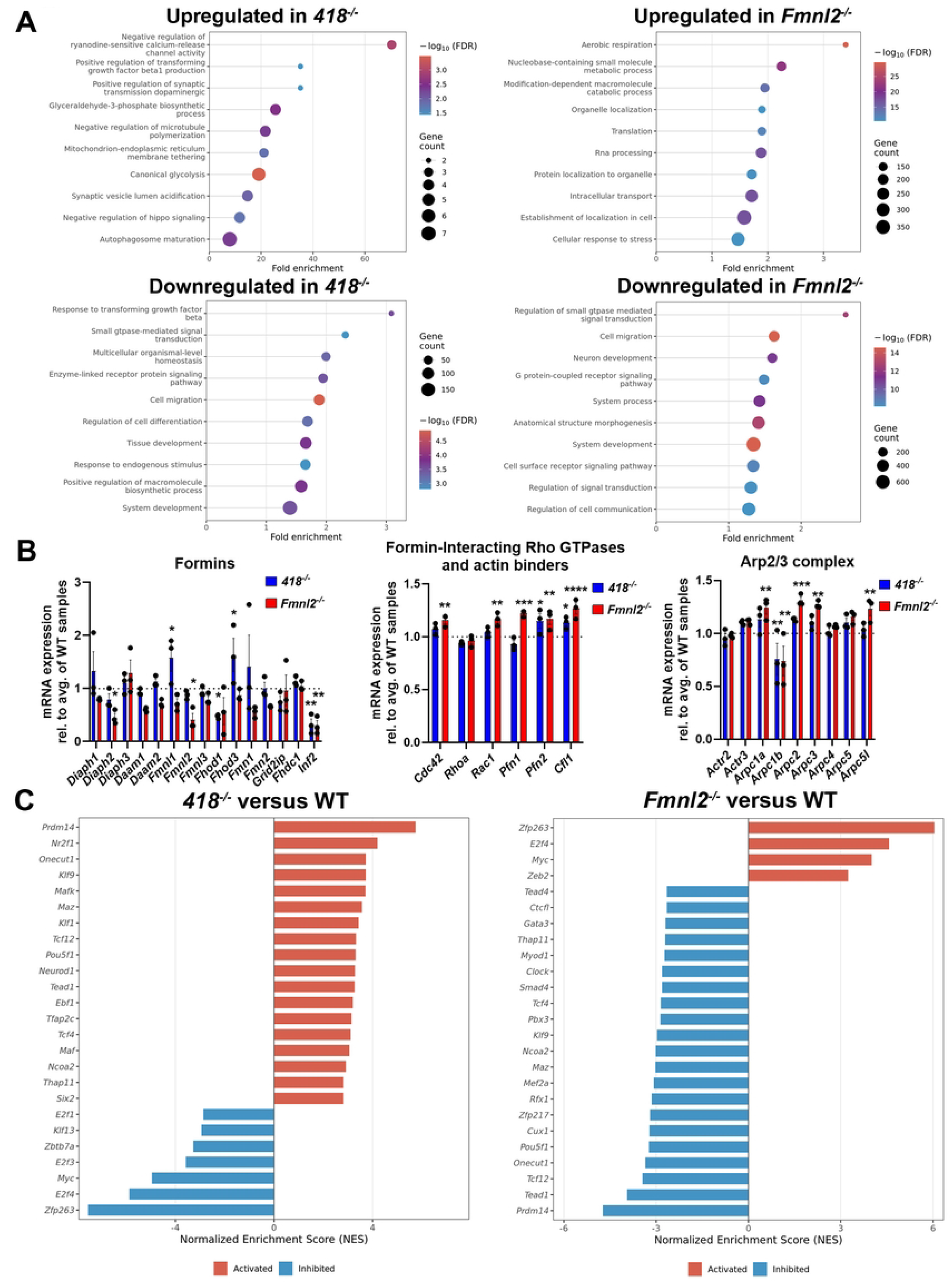
Transcriptomic analysis distinguishes hyperactive and null *Fmnl2* alleles *in vivo*. **A.** Charts depicting non-redundant gene ontology (GO) terms, identified via ShinyGO [118] that are significantly enriched among up- and downregulated in *nmf418^-/-^* and *Fmnl2^-/-^* cerebella compared to WT. **B.** Expression of formin and actin-related genes in *nmf418^-/-^* and *Fmnl2^-/-^* cerebella relative to the average of WT samples (depicted as the dashed bar at y = 1). **C.** Normalized enrichment score plots generated with DoRothEA depicting transcription factors whose activity is inferred to be activated (red) or inhibited (blue) based on differential expression analysis. Notice that several transcription factors exhibit opposite predicted activity changes in *Fmnl2^418/418^* and *Fmnl2^-/-^* cerebella. Significance was calculated by two-way ANOVA with Dunnett’s multiple comparisons test. * = p < 0.05; ** = p < 0.01; *** = p < 0.001; **** = p < 0.0001.

The upregulation of transcripts encoding glycolytic enzymes in *Fmnl2^418/418^* cerebella as shown by GO analysis may reflect a metabolic response to ionic dysregulation. Voltage-gated calcium channel positioning has been linked to the regulation of glycolysis through phosphofructokinase [70], a rate-limiting enzyme of this pathway, and actin dynamics contribute to the anchoring of glycolytic enzymes at sites of high energy demand [71]. Together, these transcriptional changes suggest that *Fmnl2*^418^ may broadly disrupt calcium homeostasis and metabolic regulation in cerebellar neurons, in addition to cerebellar morphology. In contrast to the glycolysis-related DEGs upregulated in the *Fmnl2^418/418^* cerebellum, upregulated DEGs observed in the *Fmnl2*^-/-^ cerebellum were enriched for GO terms relating to aerobic respiration. Previous reports have linked actin polymerization by INF2 and FMNL1 to mitochondrial fission and fragmentation [72–74], emphasizing a connection between F-actin formation and mitochondrial function. Interestingly, FMNL2 drives proper ER and mitochondrial spacing during development[75], suggesting that its deletion may disrupt normal organelle distribution and alter downstream mitochondrial function. The enrichment of nucleobase metabolism among upregulated genes in *Fmnl2^-/-^* cerebella may derive from the endolysosome-mTORC1-nucleotide synthesis axis, wherein actin structures at the endolysosome promote mTORC1 activity [76] which in turn drives pyrimidine synthesis [77]. Ultimately, aerobic respiration and nucleotide synthesis are intimately linked [78], which may explain their joint high enrichment.

Genes that were downregulated in *Fmnl2^-/-^* cerebella are largely related to neuronal development, small GTPase function, and cell migration, even though there is a lack of overt cerebellar phenotypes in these mice. We therefore examined whether loss of *Fmnl2* triggers compensatory changes in actin regulatory pathways that could account for this phenotypic resilience. The formins *Diaph2* and *Fmnl2*, were specifically downregulated in *Fmnl2^-/-^* cerebella, while *Inf2* was downregulated in both *Fmnl2^-/-^* and *Fmnl2^418/418^* samples, suggesting that loss of *Fmnl2* may trigger negative feedback reducing expression of other formins (Figure 5B). In contrast, mRNAs encoding the Rho GTPases CDC42 and RAC1, the actin-binding proteins profilin 1 (PFN1) and profilin 2 (PFN2), which bind actin monomers and promote their incorporation into filaments [79], and cofilin 1(Cfl1), the predominant cofilin in the cerebellum [80], which severs actin filaments and promotes filament turnover, were upregulated specifically in *Fmnl2^-/-^* but not *Fmnl2^418/418^* cerebella (Figure 5B). Several members of the Arp2/3 complex, which nucleates branched actin filaments [81], were also upregulated specifically in the *Fmnl2^-/-^* cerebella. Together, these coordinated changes across multiple actin regulatory pathways suggest that *Fmnl2*^-/-^ cerebellar cells undergo broad transcriptional remodeling to maintain actin filament homeostasis.

To identify transcription factors potentially driving these DEGs, we used DoRothEA [82, 83] and Viper [84] (Fig 5C, Table S1). Together, these programs infer relative transcription factor activity based on transcriptional changes of respective transcription factor target genes. Several transcription factors showed higher inferred activity in opposite directions between genotypes, e.g., higher in *Fmnl2^-/-^* lower in *Fmnl2^418/418^* mice including MYC, ZFP263, and E2F4. MYC is a downstream effector of Sonic Hedgehog signaling, which drives the developmental expansion of cerebellar granule cells [85–87]. The inferred downregulation of MYC activity in *Fmnl2^418/418^* cerebella may therefore contribute to cerebellar hypoplasia observed in these mice. E2F4 and ZFP263 respectively promote the specification of neural structures and the transition from pluripotency to lineage specification [88, 89]. Their reduced activity in the *Fmnl2^418/418^* cerebellum may likewise contribute to the neural phenotypes in these mice. Notably, inferred E2F4 and ZFP263 activity is elevated in *Fmnl2^-/-^* mice but not associated with an overt cerebellar phenotype. Ultimately, this DoRothEA analysis reveals discordant transcription factor activities for *Fmnl2^-/-^* and *Fmnl2^418/418^* cerebella and is consistent with our conclusion that the *Fmnl2^418^* allele acts through a gain-of-function mechanism.

## Discussion

Formins are central regulators of cellular function through their role in actin dynamics [90]. Both loss- and gain-of-function mutations in genes encoding these proteins have been linked to human disease [33–37]. Using forward genetics, we identified the causative mutation in *nmf418* mice, an ENU-induced strain with severe hindlimb gait defects, as a splice acceptor site mutation in *Fmnl2* that causes skipping of exon 10. *Fmnl2^418/418^* embryos and surviving postnatal mice have a large, skin-covered opening in the spinal column reminiscent of spina bifida occulta [61], resulting from abnormal development of the vertebral column. Mutant mice also have hydrocephalus, loss of corpus callosal axons, and abnormal cerebellar and midbrain morphology. The requirement of homozygosity paired with a dramatic reduction in FMNL2 protein abundance initially suggested a loss-of-function mechanism. However, this hypothesis was inconsistent with the absence of spinal column and brain defects in knockout mice that completely lack *Fmnl2* expression.

Exon 10 of *Fmnl2* encodes 25 amino acids within the DID autoregulatory region. Previous studies have demonstrated that mutations in the DID of other DRFs can compromise formin autoinhibition thus increasing formin activity [91, 92]. Consistent with this, our cell-based assays demonstrated that FMNL2Δ25 has increased actin polymerization activity relative to wild-type FMNL2 as measured by F:G actin fractionation and SRE-luciferase reporter assays, supporting the conclusion that loss of exon 10 produces a hyperactive protein. The functional consequence of this hyperactivity was further evident in wound healing assays, in which *Fmnl2^418/418^* MEFs showed increased migration relative to wild-type cells, in contrast to *Fmnl2*^-/-^ MEFs which showed reduced migration. Transcriptomic analysis also revealed a clear distinction between the changes resulting from the *Fmnl2^418^* mutation and loss of *Fmnl2*, demonstrating that the downstream impacts of the *nmf418* mutation diverge from those in the knockout.

The reduced endogenous levels of FMNL2Δ25 observed in *Fmnl2^418/418^* tissues are likely a consequence of its hyperactivity rather than a cause of the phenotype. A similar relationship between hyperactivity and reduced protein abundance has been described for the gain-of-function INF2 R218Q mutation, where increased actin regulatory activity is accompanied by reduced endogenous protein levels [91]. Mechanistically, the DID/DAD interaction has been shown to stabilize DIAPH3, whereas disruption of this interaction in a hyperactive DIAPH3 mutant leads to destabilization of the C-terminal DAD-containing region via the E3 ubiquitin ligase STUB1 [53]. Our co-transfection experiments revealed analogous destabilization of the C-terminal region of FMNL2 upon loss of exon 10, suggesting that FMNL2Δ25 similarly undergoes proteasomal degradation, accounting for its reduced abundance in mutant tissues. The dramatic reduction in FMNL2Δ25 levels likely underlie the recessive inheritance of this mutation and consistent with this, compound heterozygous mice with a *Fmnl2^418^* allele and a knockout allele do not display the mutant phenotypes, indicating that a minimum threshold of FMNL2 Δ25 protein is required to produce the observed abnormalities.

*Fmnl2* is broadly expressed with particularly high expression in neural tissues. However, as observed with other hyperactive formin mutations [42–46, 91], *Fmnl2^418/418^* mice display highly specific phenotypes. This specificity is likely determined not only by the tissue distribution of hyperactive FMNL2, but also by the differential sensitivity of developmental processes to aberrant actin dynamics. Many aspects of brain development rely on tightly regulated actin dynamics [93–95]. During neurogenesis, the actin network influences neurogenesis by controlling progenitor polarity and division [93]. FMNL2Δ25 may therefore perturb neurogenesis in the superior and inferior colliculi which are enlarged in *Fmnl2^418^* mutant mice and in the cerebellum, which is correspondingly decreased in size. Interestingly, brain-specific deletion of the cytoskeleton-associated Rho GTPase Rac1 affects these structures in a similar fashion [96]. FMNL2 is also an important effector of signaling molecule trafficking. PKCα-mediated phosphorylation of FMNL2 within its DAD region releases its autoinhibition and triggers its relocation to endosomes where it regulates the endocytosis of B1-integrin and cell invasion [97]. Recently, FMNL2 has been shown to associate with the trans-medial Golgi compartment via Cdc42-dependent activation, and combined knockout of *Fmnl2* and *Fmnl3* altered trafficking from the Golgi to the plasma membrane [98]. This suggests that the GTPase-independent FMNL2Δ25 activity could lead to mistimed or excessive trafficking of signaling molecules, non-cell autonomously disrupting midbrain-hindbrain patterning, as observed in mice overexpressing the homeobox transcription factor OTX2 [99]. Additional studies using mice conditionally expressing FMNL2Δ25 will be necessary to test this hypothesis.

Aberrant actin dynamics may also contribute to the hydrocephalus observed in *Fmnl2^418/418^* mice through effects on non-neuronal cells in the brain. Multi-ciliated ependymal cells line the cerebral ventricles and contribute to cerebrospinal fluid movement and defects in ependymal ciliary structure, motility, or planar polarity can disrupt ventricular fluid dynamics and have been associated with hydrocephalus [100–103]. Proper organization and stability of basal bodies, the centriole-derived structures from which ciliary axonemes extend, are required for coordinated cilia beating. In ependymal cells, the apical actin network surrounding basal bodies contributes to their stability and this network is sensitive to formin inhibition [104]. Hyperactive FMNL2Δ25 could therefore promote excessive or spatially misregulated actin polymerization in ependymal cells, disrupting basal body organization or ciliary function, contributing to the ventricular enlargement observed in *Fmnl2^418/418^* mice.

*Fmnl2^418/418^* mice have postnatal degeneration of the corpus callosal axons. Degeneration of these axons may reflect both cell-autonomous effects of hyperactive FMNL2Δ25 on the actin cytoskeleton and non-cell autonomous consequences of ventricular enlargement. The axonal actin network plays an active role in maintaining axon integrity, and mechanical injury or metabolic stress can cause actin disassembly, leading to axon fragmentation and loss of structural integrity [105]. However, hydrocephalus may also contribute to the degeneration of corpus callosal axons [106–108]. Ventricular enlargement can mechanically stretch or compress periventricular axon tracts, including the corpus callosum, and may reduce regional blood flow resulting in ischemia or metabolic stress[108]. In addition, hydrocephalus can induce neuroinflammation, which has been associated with corpus callosum injury, and reduced myelination [109–111].

The vertebral defects in *Fmnl2^418/418^* mice are reminiscent of those observed in mouse models with disruptions in the PDGF/PDGFRα pathway which regulates the migration of the dorsal sclerotome, the progenitor population that give rises to the vertebrae dorsal spinous processes [112]. PDGF/PDGFRα drives the temporally and spatially controlled activation of rho GTPases Rac1 and Cdc42 by activating phosphatidylinositol 3-kinase (PI3K). Expression of FMNL2Δ25 in the dorsal sclerotome could override this spatiotemporal control, driving GTPase-independent aberrant cell migration and the failure of spinous process formation. Alternatively, expression of this protein could disrupt the secretion of migratory cues or PI3K activators required for normal sclerotome migration, as discussed above.

While loss or reduction of FMNL2 in cell culture models have been shown to decrease filopodia number and growth [113, 114] and suppress cell motility - as we also observed in *Fmnl2*^-/-^ MEFs [115], *Fmnl2*^-/-^ mice exhibit no gross neural or anatomical phenotypes. This absence of phenotypes may reflect compensatory gene regulation changes that restore actin dynamics. Transcriptomic analysis supports this possibility and revealed upregulation of key genes several Rho GTPases, actin binders, and members of the Arp2/3 complex, the latter mirroring the observation that acute knockdown of *Arp3* induces upregulation of both FMNL2 and FMNL3 [116]. However, our analysis of *Fmnl2*^-/-^ mice did not address whether loss of *Fmnl2* affects phenotypes under specific environmental or stress conditions. For example, shRNA-mediated depletion of *Fmnl2* in tumor cells reduces tumor formation in mice [115], while morpholino-based depletion in a zebrafish model of amyloidosis reduced gliovascular remodeling, microglial activation, and amyloidβ accumulation [117]. Whether constitutive loss of *Fmnl2* similarly affects cancer progression, amyloidosis, and other disease or stress contexts awaits to be determined.

Formins are essential regulators of cytoskeletal dynamics across a broad range of developmental processes. Our investigation of gain-of-function and knockout *Fmnl2* mutant mice demonstrates that vertebrate development is exquisitely sensitive to the autoinhibitory regulation of FMNL2 and underscores the necessity of mechanistic assessment of gene variants, distinguishing loss-from gain-of-function effects to fully understand their contribution to disease pathogenesis.

## Materials and Methods

### Mice

The *nmf418* mutation was induced by N-ethyl-N-nitrosourea (ENU) mutagenesis of C57BL6/J (B6J) male mice. These mice were crossed to BALB/cByJ (BALB) female mice and G1 males were mated to BALB females. G2 females were backcrossed to G1 parental males to generate homozygous mutant mice. Genetic mapping using polymorphic microsatellite markers and single nucleotide polymorphisms was performed on mice from this cross and F2 mice from a B6J;BALB and NOD/ShiLtJ (NOD) cross. To generate B6J.*Fmnl2^418^* congenic mice, B6J;BALB-*Fmnl2*^418/+^ mice were backcrossed to B6J for at least 9 generations. Mice were genotyped using the following primers: *nmf418* WT forward: 5’TGATAACCACGTGTGTTTGA3’, *nmf418* mut forward: 5’TGATAACCACGTGTGTTTGC3’, *nmf418* reverse: 5’CTAACCTCGTTTG CCTGAGC3’.

To generate mice carrying the intron 9 A>C SNP identified in *nmf418* mice, Cas9 mRNA, a guide RNA targeting *Fmnl2* (5’CCACGTGTGTTTCAGGTCTGCGGGGTGCACACAAAGTCCAGACGCCCCGCAGACCTGAAACACACGTGG3’), and a donor DNA template introducing the single nucleotide mutation (5’TGTAATGTCTTATGCATGATAACCACGTGTGTTTCCGGTCTGCGGAGAAAAGCAGCGTTTTGAGAAGCT3’) were injected into the pronucleus of C57BL/6N (B6N) embryos. Injected embryos were implanted into pseudo-pregnant females, and the resulting offspring were genotyped using the primers above. To confirm the absence of additional mutations in this region, 400-500bp upstream and downstream of the introduced mutation was Sanger sequenced. Founder mice were crossed to B6J for at least 3 generations before intercrossing to produce homozygous *Fmnl2*^In9A>C/In9A>C^ mice.

Conditional *Fmnl2* knockout mice were generated by pronuclear injection of Cas9 mRNA, three guide RNAs (5’TAGGGGAAATAATCTCCTTT3’, 5’TTAGGATATTACATAAAGTA3’, 5’TCAGATAATCCTTGGTCCTT3’), and donor DNA oligo (5’GCTAAACAAATAGGCATAATGGTCATTATTATAGTATTTGTGGGTAGGGGAAATAATCTCCTTTAGGATATTACATAAAGTAAGTAATTCTCCTTGTCCTTTCTTCTAAGTTATATTTCACAACCAAGATTTGATAACTTCGTATAGCATACATTATACGAAGTTATAGGAAATACGATAAATATATGTTAGCATAAAGTATATTTCATATTGCATTAAATTTATTATAAATTGCAGTAGCACCAGGTACTAAGGTCATATTTTGTTGCATTATAGATATTCAGAAGGCGTGTTCAAGAATCTACACAAGTGCTAAGAGAACTGGAAATTTCATTAAGAACCAACCACATTGGGTAAGT ATGCTCCTGTAACATGTTAGCACTAACTTTTTATTTTTAGAAAGTGACTGAATCATACCTAAGAACTGAGCTTCTATATTTACCGAAGATAACTTCGTATAGCATACATTATACGAAGTTATGACCAAGGATTATCTGAGGGAAGTCTTCCTTAGTCCCCTGATCACTGACTGTGAGAGTCAGTAGAAGCAGAATGGGGAGCCCTTGACGTGGCTATTTATAGCAATTCTGTGTTTACTTGTTTGGATGAGGTAAGTACAGACAGCTCATGAC3’) into C57BL/6J embryos to insert loxP sites flanking exon 4 of *Fmnl2*. Mice containing the modified allele were identified using primers TMF1980 5’ TCATAAGTGACTCCATGGGCAA3’ and TMF1982 5’ GAAAAGCCTCTGAAGACTATGCC3’ and mated to B6.Cg-*Edil3^Tg(Sox2-cre)1Amc^*/J (Sox2-Cre) mice to generate offspring with Cre-dependent deletion of *Fmnl2* in the epiblast [119].

Mice were group-housed on a 12-hour light/12-hour dark cycle. Both sexes were used for all experiments. Embryonic samples were used without sex determination. All mouse studies were conducted in accordance with the institutional and regulatory guidelines of the University of California San Diego and The Jackson Laboratory.

### Histology and Immunofluorescence

For histological analysis, anesthetized mice were transcardially perfused with 4% paraformaldehyde (PFA) or 10% neutral buffered formalin (NBF), and brains were post-fixed overnight prior to paraffin embedding. 7μm sections were stained with hematoxylin and eosin, cresyl violet, or cresyl violet/Luxol fast blue according to standard protocols [74, 75]. Histological images were acquired using a Hamamatsu digital slide scanner. For gross anatomical imaging, brains were immersed in Bouin’s fixative overnight and imaged using a Nikon SMZ 1500 dissection microscope equipped with an RT SE SPOT camera (Diagnostic Instruments). Some representative images were acquired on a VS200 Olympus Slide Scanner.

For L1CAM immunofluorescence, tissue from embryonic mice was flash frozen in liquid nitrogen and embedded in optimal cutting temperature compound (Electron Microscopy Sciences, 62550). 30μm sections were fixed in ice-cold methanol. Sections were incubated overnight at 4°C with an antibody against L1CAM (L1, 1:250, Millipore Sigma, MAB5272) then goat anti-rat Alexa Fluor-555 (ThermoFisher, #A21434; 1:100) for three hours at room temperature. Nuclei were counterstained with DAPI. Images were acquired on a Nikon CSU-W1 SoRa spinning disk confocal microscope.

For pan-phospho neurofilament immunofluorescence, tissue from embryonic mice was fixed in 4% PFA overnight without perfusion and embedded in paraffin. Antigen retrieval was performed on 7μm sections by heating slides in 0.01M sodium citrate buffer with 0.05% Tween-20 (pH 6.0) for three cycles of three minutes in a microwave. Each cycle was followed by incubation at room temperature for three minutes. Sections were incubated overnight at 4°C with an antibody cocktail against neurofilament (SMI-312, 1:100, BioLegend #837904) then goat anti-mouse Alexa Fluor-488 (ThermoFisher, #A11029; 1:500) for two hours at room temperature. Nuclei were counterstained with DAPI and autofluorescence was quenched with Sudan Black. Images were acquired on a Zeiss Axiovert 200 microscope.

### RT-PCR and qPCR

Cerebellar tissue from P21 WT and *nmf418^-/-^* euthanized mice was flash-frozen in liquid nitrogen and stored at -80°C. RNA was isolated by using TRIzol (Life Technologies, #15596018). RNA concentration and purity (A260/A280) were assessed by NanoDrop. cDNA was synthesized from DNase-treated total RNA using oligo(dT) primers with the SuperScript III or IV First-Strand Synthesis System (ThermoFisher, #18080051, #18090200). RT-PCR was performed for 30 cycles using the following primers: exon 7 forward: 5’-TTGCCAAGCAGAAGAACCCT-3’; exon 10 reverse: 5’-TCAGCTTCTCAAAACGCTGC-3’; exon 9 forward: 5’-CAGCCGTTTGTCTTGTCAGA-3’; exon 12 reverse: 5’-CGCTCTCTGTGTGTTTCAGC-3’. Quantitative RT-PCR was performed using 40ng of cDNA with 2-3 technical replicates using SYBR Green Supermix (#1708880, Biorad) with the following primers: exon 13 forward: 5’ TTGTCCACGGGGTCAGAACTA 3’, exon 14 reverse: 5’ GTGGCGATGCTGTTCCATT 3’, exon 9 forward: 5-TTGTCTTGGAGCTGTTGGCA-3’, exon 10 reverse: 5’-AAAACGCTGCTTTTCTCCGC-3’, GAPDH forward: 5’-AGGTCGGTGTGAACGGATTTG-3’, GAPDH reverse: 5’-TGTAGACCATGTAGTTGAGGTCA-3’ [120]. Technical replicates were averaged to obtain the Ct values for each sample, and relative transcript abundance was calculated using the ΔΔC_t_ method [121]. Statistical significance was determined by unpaired T-test using GraphPad Prism 10.

### Western Blots

P21 tissues and mouse embryonic fibroblasts were flash-frozen in liquid nitrogen and homogenized in RIPA buffer (150mM NaCl, 1% NP-40, 0.5% Deoxycholic acid, 0.1% SDS, 50mM Tris pH 7.4) containing protease inhibitors (Sigma, #11836170001). Total protein concentration was determined by DC Protein Assay (Bio-Rad, #5000112). Proteins were resolved on SDS/PAGE gels and transferred to PVDF membranes (Amersham, #1060023). Western blots were performed using standard methods with the following antibodies: rabbit anti-FMNL2 (1:1000, HPA005464, rabbit, Sigma, #HPA005464), anti-Hsp90 (1:1000, Hsp90D7-alpha, rat, Stressgen, #Hsp90D7α), anti-GFP (1:5000, mouse, Chemicon, MAB3580), anti-RFP (1:5000, mouse, Advanced Targeting Systems, AB-332), anti-β-tubulin (1:2000, rabbit, Cell Signaling Technologies, 2146), anti-FLAG (1:5000, rabbit, Cell Signaling Technologies, 14793), anti-Myc tag (1:10000, goat, Abcam, 9132), anti-actin (1:10000, rabbit, Cell Signaling Technologies, 4970), HRP-conjugated secondary antibodies used were goat anti-rabbit (1:2000, Bio-Rad, #1706515), anti-rat (1:2000, Bio-Rad, #5204-2504), anti-mouse (1:5000, Bio-Rad, #1706516), and donkey anti-goat (1:5000, Jackson ImmunoResearch, 705-035-003). Signal was detected with SuperSignal West Pico PLUS chemiluminescent substrate (ThermoFisher, 34580) on film (BioPioneer, MUBF-01) or Azure Biosystems 600 western imager.

### Skeletal stains

E18.5 embryos were euthanized, skin and fascia were removed, and carcasses were fixed in 95% ethanol overnight at room temperature. Ethanol was replaced with acetone and incubated overnight at room temperature. Cartilage was stained overnight in 0.03% Alcian blue in 20% glacial acetic acid/80% ethanol, followed by two brief washes in 70% ethanol and overnight incubation in 95% ethanol. Tissue was cleared in 1% KOH for 1 hour, then bone was stained overnight in 0.005% Alizarin red in 1% KOH at 4°C. Specimens were destained by incubation in 50% glycerol/1% KOH over several days and stored in 100% glycerol. Images were acquired on a Nikon SMZ 1500 dissection microscope equipped with an RT SE SPOT camera (Diagnostic Instruments).

### RNA Sequencing Library Preparation

E18.5 cerebella were flash-frozen in liquid nitrogen and homogenized in 500 µL TRIzol (Life Technologies, #15596018). Total RNA was isolated according to the manufacturer’s protocol. On-column DNase digestion and RNA purification were performed using the RNeasy Mini Kit (Qiagen, #74104), RNA quality was assessed on an Agilent TapeStation. Libraries were prepared using the SMART-Seq mRNA LP Kit (Takara Bio, #634769) according to the manufacturer’s protocol. Library concentration was quantified on a Qubit 4 fluorometer, and average fragment length was determined on the Agilent TapeStation. Paired-end reads (2 × 100 bp) were obtained on an Illumina NovaSeq X Plus platform.

### RNA Sequencing Analysis

Raw paired-end FASTQ files were quality-trimmed using fastp (v0.23) [122] with automatic adapter detection, a minimum read length of 36 bp, a minimum base quality of Q20, and paired-end error correction enabled. Trimmed reads were pseudoaligned to the GRCm39 mouse transcriptome (Gencode release M36) using Salmon [123] with GC bias correction (--gcBias), sequence-context bias correction (--seqBias), and mapping validation (--validateMappings). Transcript-level abundances were imported and summarized to gene level using tximport with the lengthScaledTPM method, retaining only protein-coding genes.

Gene-level counts were analyzed with DESeq2 [124]. Genes with fewer than 10 total counts across all samples were excluded prior to model fitting. Two pairwise contrasts were evaluated: *Fmnl2*^418/418^ vs. WT and *Fmnl2*^-/-^ vs. WT, each using apeglm log₂ fold-change shrinkage. Differentially expressed genes were defined by an adjusted p-value < 0.05 and |log₂FC| ≥ 1. One WT replicate was excluded from all analyses after principal component analysis identified it as a clear outlier.

Per-event percent-spliced-in (PSI) values were calculated from Salmon TPM output using SUPPA2 [125], considering five event types: skipped exons (SE), alternative 5′ and 3′ splice sites (A5SS/A3SS), mutually exclusive exons (MX), retained introns (RI), and alternative first and last exons (AF/AL). Differential splicing between *Fmnl2*^418/418^ and WT was assessed using the SUPPA2 empirical method. For direct quantification of reads mapping to individual *Fmnl2* exons, trimmed reads were aligned to Chromosome 2 of GRCm39 using HISAT2 [126] with splice-site-aware parameters. Alignments were coordinate-sorted with SAMtools [127]. Exon-level read counts were obtained using featureCounts [128] at exon resolution (-f -p -B -- fracOverlap 0.5 -s 2), and coverage was normalized within each sample to counts per million and expressed relative to the maximally covered exon. Splice junction read counts and per-base coverage were extracted from exon-filtered BAM files using GenomicAlignments [129] and visualized as sashimi plots.

Transcription factor (TF) activity was inferred from the DESeq2 differential expression results using the DoRothEA [82, 83] regulon and the VIPER [84] algorithm. The DoRothEA mouse regulon was retrieved from the dorothea R package and converted to VIPER format using the df2regulon function. For each contrast, DESeq2 Wald test statistics were used as a gene-level ranking metric and assembled into a gene by contrast matrix. TF activity scores were computed using the viper function yielding a normalized enrichment score (NES) for each TF in each contrast, where positive NES indicates inferred activation and negative NES indicates inferred inhibition relative to WT.

### CAD Cell Overexpression and Imaging

Cath.a-differentiated (CAD) cells were originally obtained from Sigma-Aldrich (#08100805). Cells were cultured in DMEM/F12 medium with L-glutamine (Gibco, #11320-033) supplemented with 8% fetal calf serum (R&D Systems, #S11150) and 1% penicillin-streptomycin (Corning, #30-002-CI). Prior to imaging, coverslips were coated overnight at 4°C with 10 µg/mL laminin (Sigma-Aldrich, #L2020) and cells were plated onto coated coverslips. DNA constructs were introduced by electroporation using the Neon Transfection System (Invitrogen, #MPK5000) with the 10 µL transfection kit. Cells were grown to 70–80% confluency, detached with 0.5% trypsin/EDTA (Corning, #25-051-CI), pelleted by brief centrifugation, and rinsed with Dulbecco’s phosphate-buffered saline (DPBS; Corning, #21-031-CV). Cell pellets were resuspended in a minimal volume of Buffer R (Invitrogen, #MPK1025) containing 1.3 µg of DNA per electroporation reaction (1 µg for mCherry-C1 control plasmid, #632523). Electroporated cells were immediately seeded onto laminin-coated coverslips and cultured overnight in antibiotic-free medium prior to imaging.

Images were acquired on a Nikon CSU-W1 SoRa spinning disk confocal microscope equipped with a 100× 1.49 NA SR objective and a Hamamatsu Fusion BT camera operating in W1 spinning disk mode. Confocal z-stacks underwent background subtraction using Denoise.ai (NIS-Elements), a deep learning–based denoising algorithm, followed by automatic 3D deconvolution (NIS-Elements). At least 20 random fields of view were acquired per condition per biological replicate.

All CAD cell data were tested for normality using the Shapiro-Wilk test. Normally distributed data were compared using a one-way ANOVA followed by Dunnett’s post-hoc test for comparisons against a single control condition or Tukey’s post-hoc test for all pairwise comparisons. Data failing the normality test were compared using the Kruskal-Wallis test followed by Dunn’s multiple comparisons test. All statistical analyses and graphing were performed in GraphPad Prism 10.

### Mouse embryonic fibroblast wound healing and proliferation assays

Mouse embryonic fibroblasts (MEFs) were derived from E12.5 embryos according to standard protocols [130]. MEFs were maintained in Dulbecco’s modified Eagle’s medium (DMEM; Gibco, #41965039) supplemented with 1× GlutaMAX (Gibco, #35050061), 1× PSN (Gibco, #15640055), 1% penicillin-streptomycin-neomycin (Invitrogen, 15640055) and 10% embryonic stem cell-qualified fetal bovine serum (Gibco, #10439024) at 37°C in 5% CO2. Cells were grown to 95% confluency and passaged at least once prior to experimentation.

For wound healing assays, passage 1 (P1) MEFs were plated in triplicate at 75% confluency in 96-well clear-bottom plates (Corning, #3997) and incubated for at least 8 hours at 37°C in 5% CO2. Wounds were introduced with a single pass of the 96-well Incucyte WoundMaker tool (Sartorius), after which the media was replaced. Wound closure was monitored by live-cell imaging for 36 hours on a Sartorius Incucyte S3 Live-Cell Analysis System, acquiring two images per well at 10× magnification every hour. Images were analyzed using Incucyte S3 software. Confluence and wound masks were generated from one representative well per condition at 0 hours and applied to all remaining wells across the 36-hour time course. Measurements that were not accurately detected by the imaging algorithm were excluded from analysis. Differences in wound closure between genotypes were assessed by two-way ANOVA with multiple comparisons.

For BrdU incorporation assays, P1 MEFs were plated to 50% confluency in glass bottom 24-well plates (CellVis P24-1.5P) and incubated with 10μM BrdU (Sigma Aldrich, B9285) for one hour prior to fixation with 4% PFA for 15 minutes. Fixed cells were permeabilized with 0.3% Triton X-100 in PBS for 15 minutes, rinsed twice with PBS, then treated with 1N HCl for 15 minutes to unmask incorporated BrdU. Cells were neutralized with 0.1M sodium borate buffer (pH 8.5) for 15 minutes and rinsed three times with PBS. Cells were blocked with 10% normal goat serum (Jackson ImmunoResearch, 005-000-121) in 0.3% PBS Triton X-100 (blocking solution) for 1 hour. Cells were incubated overnight at 4°C with mouse anti-BrdU (Abcam, ab8955, 1:500) and rabbit anti-KI67 (Abcam, ab15580, 1:500) diluted in blocking solution. After three 5-minute washes in 0.3% PBS Triton X-100, cells were incubated with secondary antibodies (Thermo Fisher Scientific, A-11029, goat anti-mouse 488, 1:1000; Thermo Fisher Scientific, A-21428, goat anti-rabbit 555, 1:1000) and DAPI (2μg/mL) diluted in blocking solution for one hour at room temperature. Cells were washed three times in PBS for 5 minutes then covered with 50% glycerol in PBS for imaging on a Nikon CSU-W1 SoRa spinning disk confocal microscope. Cells positive for BrdU and/or KI67 were counted across three fields of view from each well and expressed as a percentage of the total number of cells. Differences between groups was determined by two-way ANOVA with Tukey’s multiple comparisons test.

### HEK293T cell culture

HEK293T cells (ATCC, CRL-3216) were maintained in DMEM containing 10% heat-inactivated fetal bovine serum (Genesee Scientific, 25-514H) in a 37°C incubator with 5% CO_2_. Cells were passaged upon reaching 85% confluency. All transfections were performed using Calfectin DNA transfection reagent (SignaGen Laboratories, SL100478) using manufacturer’s protocol.

### Plasmids

WT FMNL2 and FMNL2 Δ25 were subcloned into the pEGFP-N1 backbone from plasmids custom synthesized by VectorBuilder. The CDC42-G12V plasmid was generated using point mutagenesis of wild-type CDC42 cloned into the pCMV-3Tag-1A backbone. 3xFLAG WT FMNL2 and Δ25 N-terminal fragment inserts were amplified with overlaps to CMV-3Tag-1A by PCR from WT FMNL2-eGFP and FMNL2 Δ25-eGFP plasmids, respectively, using the following primers: forward: 5’-TGACGATAAAGCCCGGGCGGAAGCTGCCGATGCCCGAG-3’; reverse: 5’-CGAGGTCGACGGTATCGATACTACTGAATTTTAATGGTCCCTTGTTTCTCC-3’. Inserts were purified by agarose gel electrophoresis and cloned into BamHI- and HindIII-digested pCMV-3Tag-1A using Gibson Assembly (NEB; E2621L), as per manufacturer’s instructions. The 3xMyc FMNL2 C-terminal fragment insert was amplified with overlaps to CMV-3Tag-2A from the WT FMNL2-eGFP plasmid using the following primers: forward: 5’-GGAAGATCTGGCCCGGGCGGCCACCACCTCAGGACCCTC-3’; reverse: 5’-CGAGGTCGACGGTATCGATACCTACCGCATCGGCTCGT-3’. The insert was purified and cloned into HindIII- and BamHI-digested pCMV-3Tag-2A as above.

### FMNL2 fragment overexpression

HEK293T cells in a 6-well plate were transfected at 50% confluence with 1μg 3xFLAG-tagged WT or Δ25 N-terminus, alone or in combination with 3xMyc-tagged C-terminus (which is the same sequence for both FMNL2 variants). Medium was exchanged 24 hours after transfection, and cells were lysed in RIPA buffer 72 hours post-transfection and analyzed via SDS-PAGE and western blot.

### Actin fractionation

HEK293T cells in a 6-well plate were transfected at 50% confluence with 2μg of plasmids encoding eGFP, WT FMNL2-eGFP, or FMNL2 Δ25-eGFP). Medium was replaced 24 hours post-transfection. 72 hours post-transfection, cells were rinsed in PBS, collected using a cell scraper and pelleted by centrifugation. Input samples were reserved for two of the three replicates prior to fractionation. G-Actin and F-actin fractions were separated using the G-Actin/F-actin In Vivo Assay Biochem Kit (Cytoskeleton, Inc., BK037) according to the manufacturer’s protocol. Resulting fractions were analyzed SDS-PAGE and western blot. Band intensities were quantified in FIJI and plotted as a ratio of F:G actin. Significance was determined by one-way ANOVA with Dunnett’s multiple comparisons test.

### Actin polymerization reporter assay

HEK293T cells in a 6-well plate were transfected at 70% confluency with 2μg total DNA (0.5μg Fluc actin reporter, 0.5μg Rluc control reporter, and 1μg eGFP, WT FMNL2-eGFP, or FMNL2 Δ25-eGFP). 24 hours post-transfection, cells were replated into white-walled, white-bottom 96-well plates at 25,000 cells per well with 6 wells per condition. After six hours, media was replaced with 100μL of starvation medium (DMEM with 0.5% FBS) for 16 hours. Media was then partially exchanged by 50μL and replacing with either control medium (DMEM) or induction medium (DMEM with 40% FBS). Cells were incubated for 6 hours. Firefly and *Renilla* luciferase activity was quantified using the Dual-Glo Luciferase Assay System (Promega, E2920) on a SpectraMax i3X plate reader. Significance was determined by two-way ANOVA with Tukey’s multiple comparisons test.

### Co-immunoprecipitation

For eGFP/mCherry co-immunoprecipitations of FMNL2 homodimers and heterodimer, HEK293T cells in a 6-well plate were transfected at 50% confluence with 1μg WT FMNL2-eGFP or FMNL2 Δ25-eGFP and 1μg WT FMNL2-mCherry or FMNL2 Δ25-mCherry. Medium was replaced 24 hours post-transfection. 72 hours post-transfection, cells were rinsed with PBS, collected using a cell scraper and pelleted by centrifugation. Pellets were lysed in IP buffer (25mM Tris, pH 7.4, 150mM NaCl, 1% NP-40, 1mM EDTA, 5% glycerol). Magnetic protein A beads per sample (50μL per sample, Invitrogen 10002D) were pre-bound with 2μL anti-GFP antibody (Invitrogen, A-11122) and incubated overnight with lysate at 4°C. Unbound fractions were collected prior to washing. Beads were washed three times with IP buffer for 10 minutes at 4°C. Input, unbound, and IP samples were analyzed by SDS-PAGE and western blot.

For FMNL2-CDC42 co-immunoprecipitation, HEK293T cells in a 10cm dish were transfected at 70% confluency with 5μg CDC42 G12V DNA with 5μg WT FMNL2-eGFP or FMNL2 Δ25-eGFP plasmids. 24 hours post-transfection, cells were rinsed with PBS, collected using a cell scraper and pelleted by centrifugation. Pellets were lysed in IP buffer. To normalize bait input (exogenous FMNL2) across immunoprecipitations, 5μL of each lysate was used for dot blot analysis with anti-GFP antibody (1:5000, mouse, Chemicon, MAB3580). To adjust for equivalent levels of WT and Δ25FMNL2 GFP-tagged proteins, lysates from WT FMNL2-eGFP-overexpressing cells were diluted with lysate from cells overexpressing CDC42 G12V alone. Normalized lysates were incubated with 100μL of magnetic protein A beads (Invitrogen 10002D) pre-bound with 5μL anti-GFP antibody (Invitrogen, A-11122) overnight at 4°C. Beads were washed three times with IP buffer for 10 minutes. Input and IP samples were analyzed by SDS-PAGE and western blot.

## Data Availability

RNA sequencing data generated in this study can be accessed with the Gene Expression Omnibus accession number GSE348874.

## Acknowledgements

RNA-Seq data were generated at the UC San Diego IGM Genomics Center utilizing an Illumina NovaSeq X Plus that was purchased with funding from a National Institutes of Health SIG grant (#S10 OD026929). Some images were acquired through the UC San Diego Microscopy Core, which is funded by NS047101. This work was supported by NIH/NIGMS R35GM137959 to E.A.V. S.L.A. is an Investigator of the Howard Hughes Medical Institute.

---

**Fig. S1.**
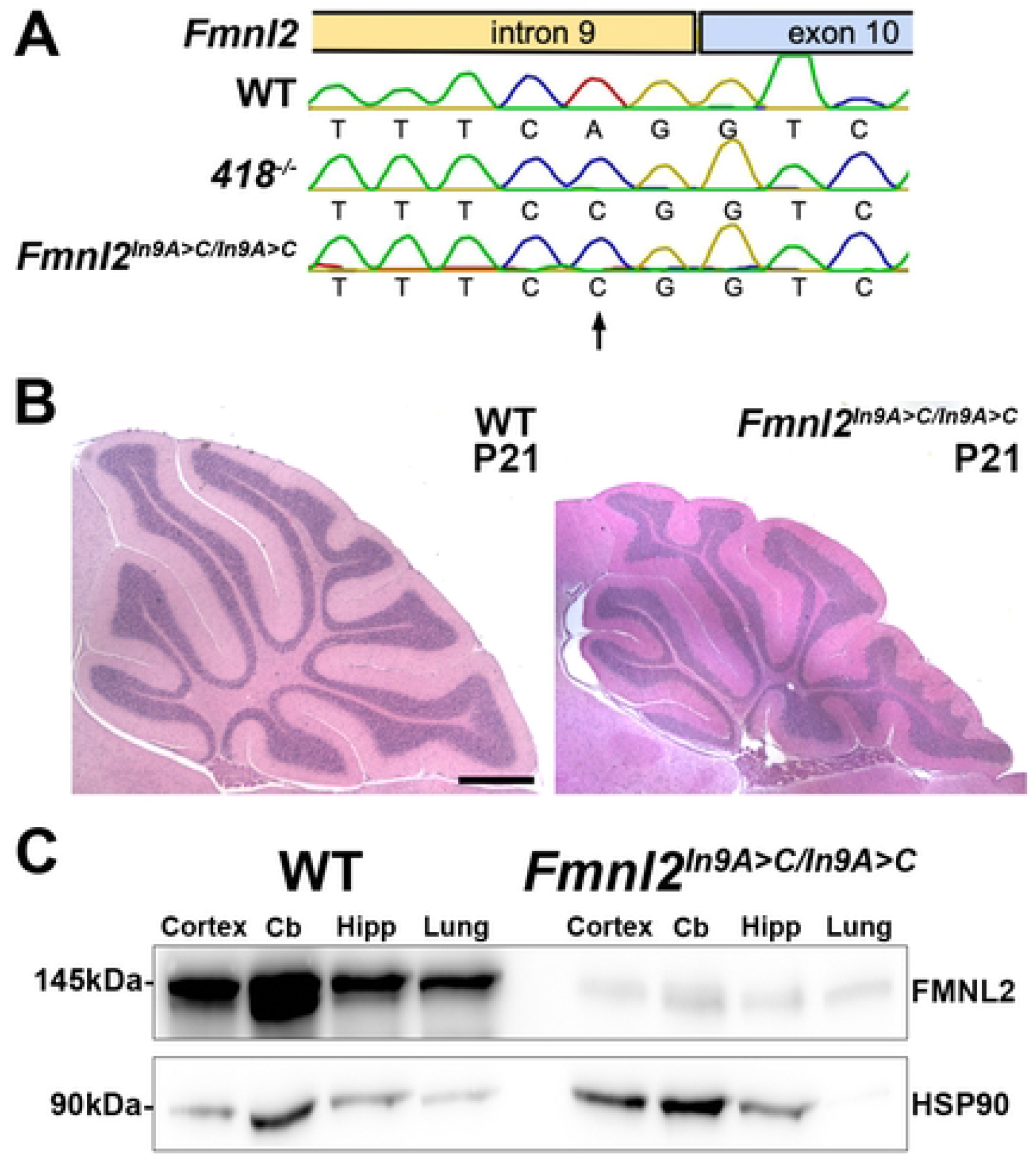
CRISPR-Cas9-mediated introduction of the *nmf418* mutation recapitulates major phenotypes. **A.** Sequencing chromatograms of the *Fmnl2* intron 9/exon 10 boundaries in wild-type (WT), *418^-/-^*, and *Fmnl2^In9A>C/In9A>C^* genomic DNA. Arrow indicates the A>C transversion at the canonical AG splice acceptor site of intron 9. The same WT and *418^-/-^* chromatograms are shown in Fig 2C. **B.** Midline sagittal sections of P21 WT and *Fmnl2^In9A>C/In9A>C^* cerebella stained with hematoxylin and eosin, demonstrating cerebellar hypoplasia and lobule disorganization in *Fmnl2^In9A>C/In9A>C^* mice. Scale bar = 200μm. **C.** Representative western blot for FMNL2 in cortex, cerebellum (Cb), hippocampus (Hipp), and lung from WT and *Fmnl2^In9A>C/In9A>C^* mice, demonstrating a dramatic reduction in FMNL2 protein levels. HSP90 was used as a loading control.

**Fig. S2.**
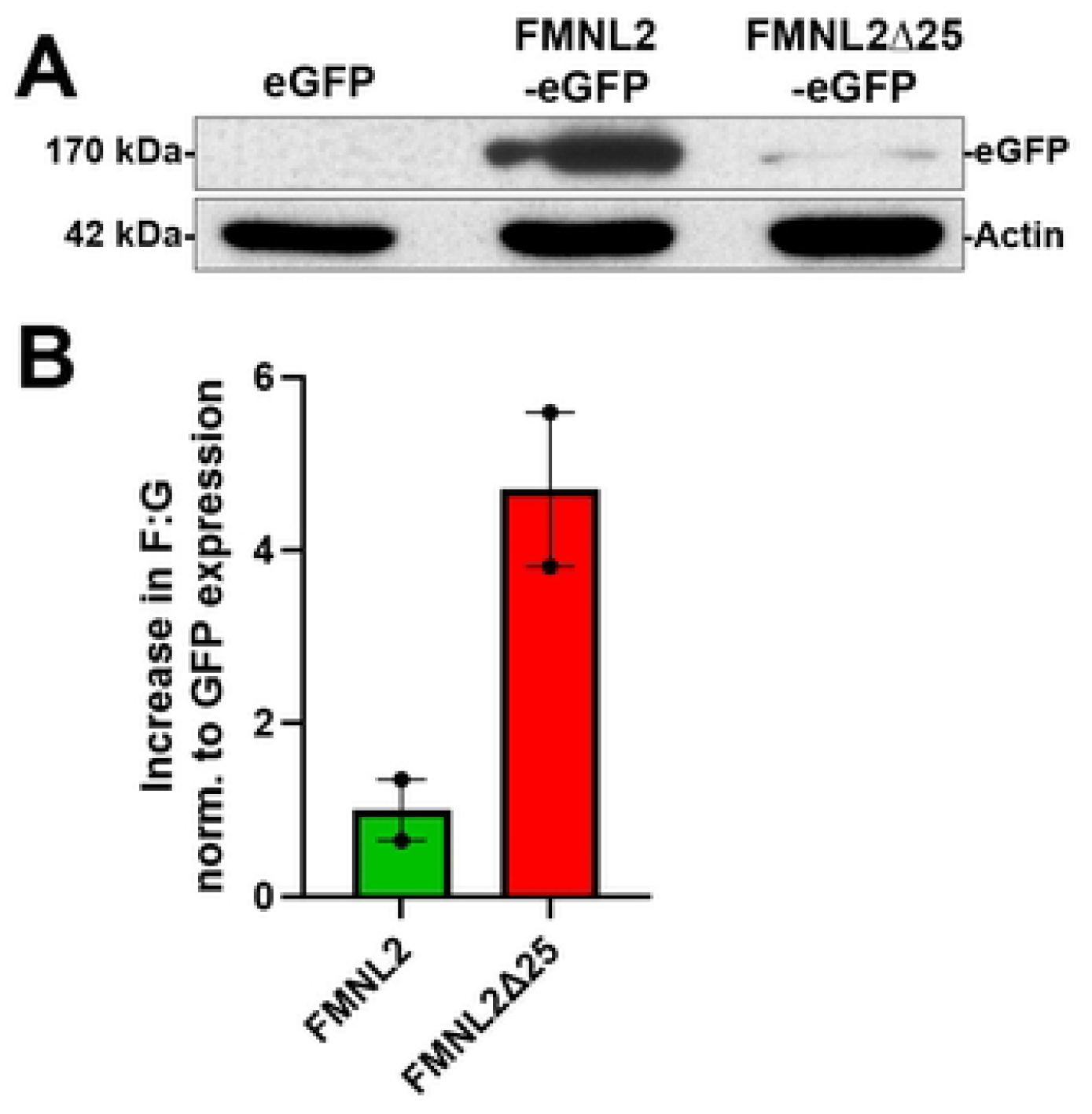
Exogenous FMNL2Δ25 promotes increased actin filamentation when accounting for protein levels. **A.** Representative western blot for eGFP and actin in input lysates from HEK293T cells (Fig. 3C, D) overexpressing eGFP, wild-type FMNL2-eGFP, or FMNL2Δ25-eGFP prior to globular (G) and filamentous (F) actin fractionation. Note the reduced expression of FMNL2Δ25-eGFP relative to wild-type FMNL2-eGFP. Input samples were available from two of the three fractionated replicates. **B.** F:G actin ratio differences from Fig. 3D normalized to the level of exogenous eGFP-tagged protein from Panel A, demonstrating an approximately 5-fold greater ratio in FMNL2Δ25-eGFP-expressing cells relative to those expressing wild-type FMNL2-eGFP.

**Fig. S3:**
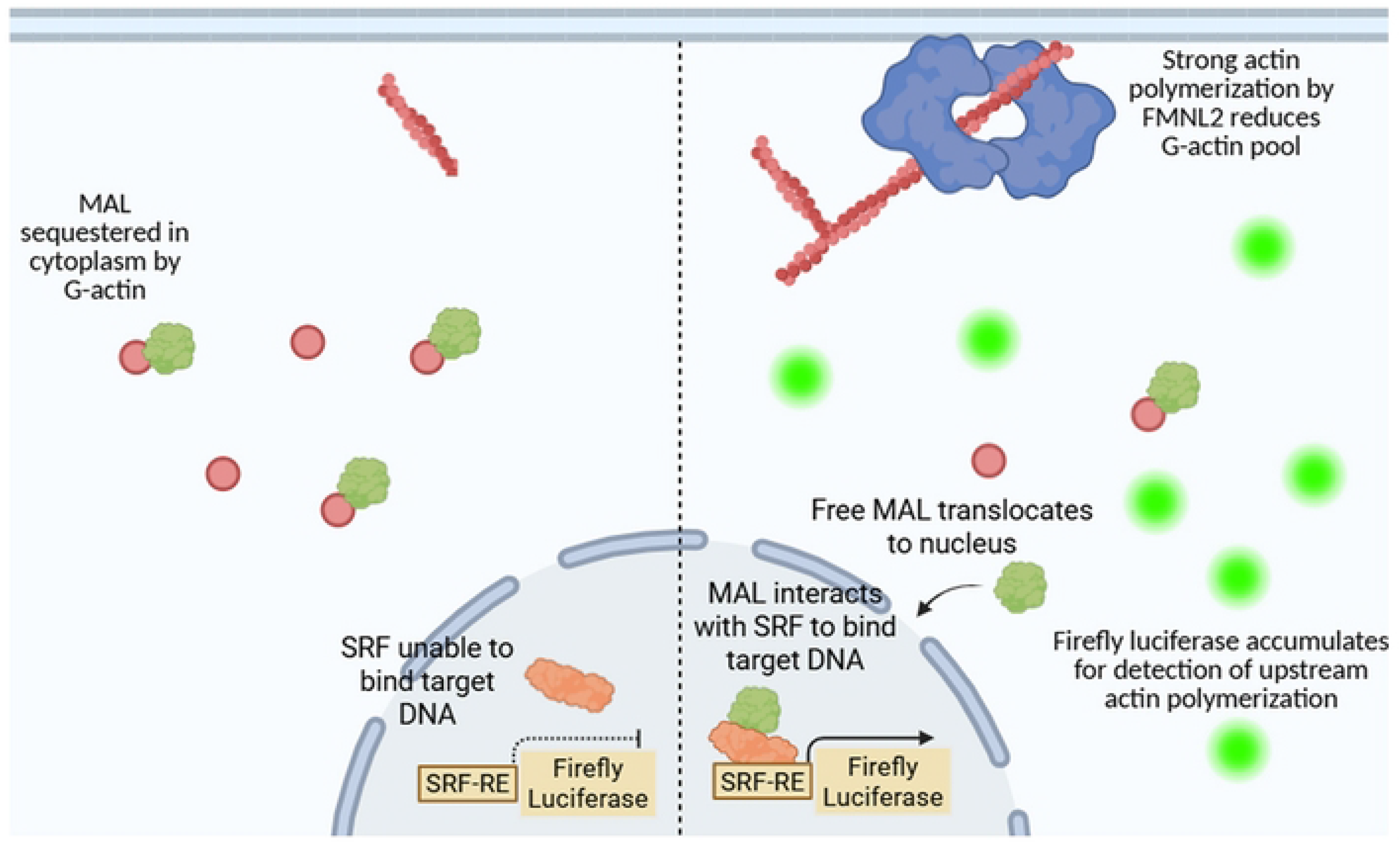
Schematic of the SRE-firefly luciferase reporter assay. Under basal conditions (left), MAL (MRTF-A) is sequestered in the cytoplasm by free G-actin, preventing serum response factor (SRF) from binding the serum response element (SRE) and driving firefly luciferase expression. Upon actin polymerization (right), the free G-actin pool is reduced, MAL translocates to the nucleus, binds SRF at the SRE, and drives firefly luciferase accumulation, providing an indirect readout of upstream actin polymerization activity.

**Fig. S4:**
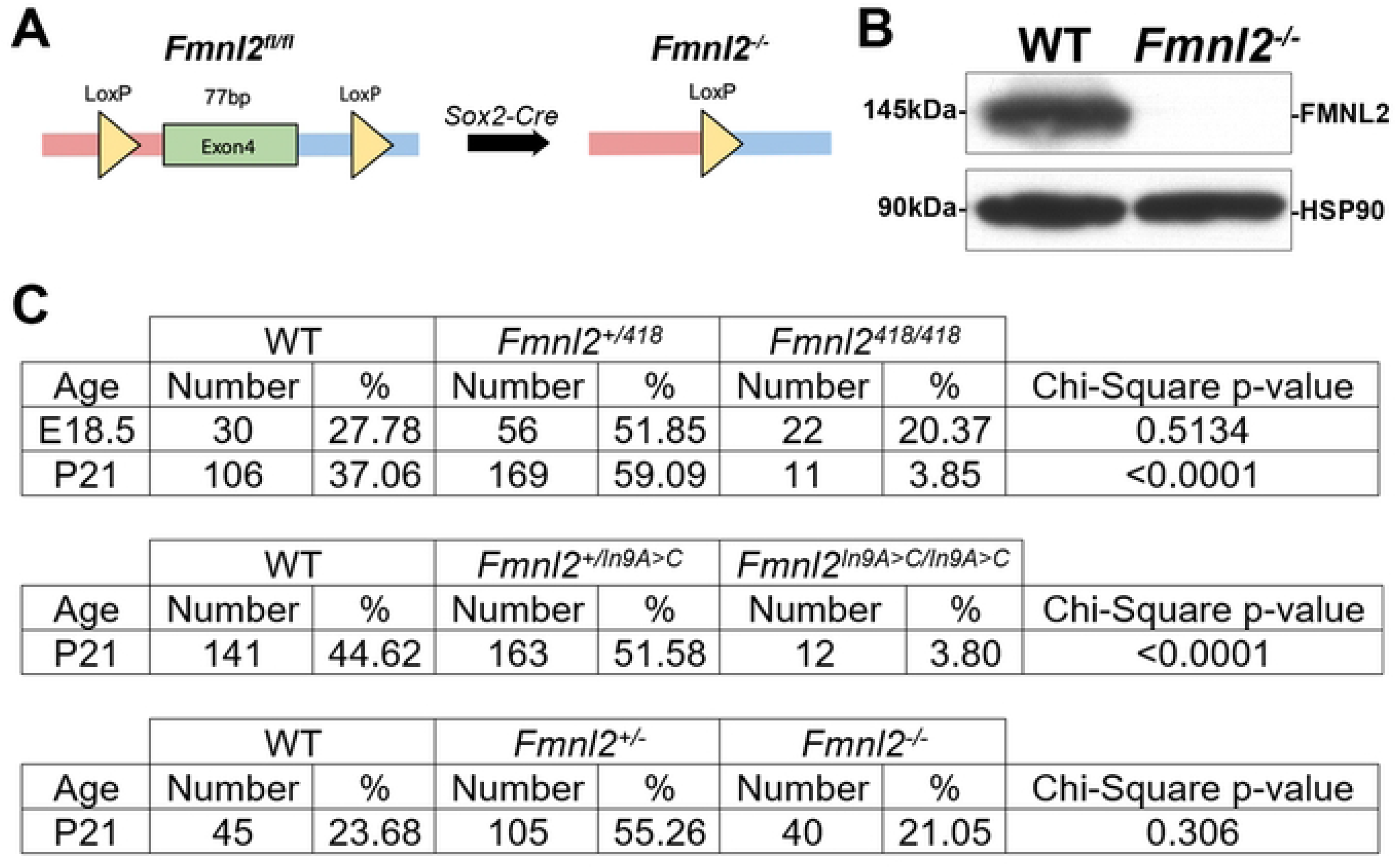
Postnatal lethality in *Fmnl2^418/418^* but not *Fmnl2^-/-^* mice. **A.** Schematic of the conditional *Fmnl2* knockout allele. LoxP sites were inserted flanking the 77-bp exon 4 to generate *Fmnl2^fl/fl^* mice. Sox2-Cre-mediated recombination deletes exon 4, causing a frameshift and premature stop codon, producing a null *Fmnl2*^-/-^ allele. **B.** Representative western blot for FMNL2 in cerebellar extracts from WT and *Fmnl2*^-/-^ mice confirming complete loss of FMNL2 protein following exon 4 deletion by Sox2-Cre. HSP90 was used as a loading control. **C.** Observed proportions of E18.5 and/or P21 progeny from intercrosses of *Fmnl2^+/418^*, *Fmnl2^+/In9A>C^*, or *Fmnl2*^+/-^ heterozygous mice. *Fmnl2^418/418^* and *Fmnl2*^I*n9A>C/In9A>C*^ homozygotes are present at Mendelian ratios at E18.5 but are underrepresented at P21 indicating postnatal lethality. *Fmnl2*^-/-^ mice are recovered at normal Mendelian ratios at P21. Statistical significance was assessed by Chi-square test.

**Fig. S5.**
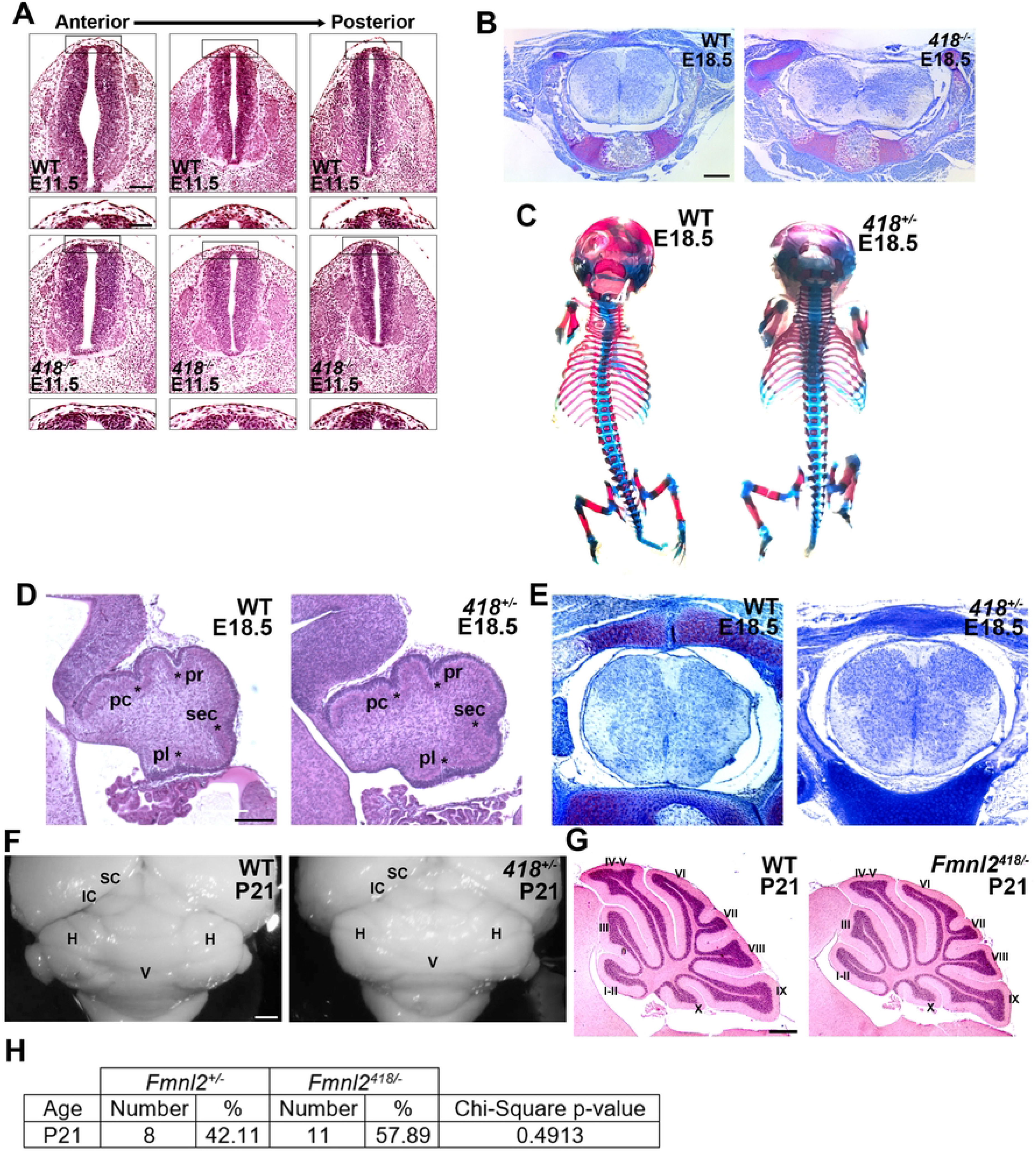
*Nmf418* phenotypes require two mutant alleles. **A.** Hematoxylin and eosin-stained coronal sections of E11.5 wild-type (WT) and *Fmnl2^418/418^* (*418*^-/-^) embryos at anterior to posterior (left to right) levels of the neural tube. Higher magnification images of the dorsal region of the neural tube (boxed) are shown below. The neural tube is closed across the entire anterior-posterior axis in *Fmnl2^418/418^* embryos, indicating that the spinal column opening is not caused by failure of neural tube closure. Top Scale bars = 100 µm (top) and 40μm (bottom). **B.** Transverse lumbar spinal sections of E18.5 WT and *Fmnl2^418/418^* mice stained with cresyl violet. Notice the flattening of the spinal cord and lack of fusion of the spinous process in the mutant. Scale bar = 200μm. **C.** Dorsal views of alcian blue (cartilage)/alizarin red (bone)-stained skeletons from WT and *Fmnl2^+/418^* (*418^+/-^*) E18.5 embryos. **D.** Midline sagittal sections of cerebella from E18.5 WT and *Fmnl2^+/418^* mice. Asterisks denote visible principal fissures. pc, preculminate fissure; pr, primary fissure; sec, secondary fissure; pl, posterolateral fissure. Scale bar = 100μm. **E.** Transverse lumbar spinal sections of E18.5 WT and *Fmnl2^+/418^* mice stained with cresyl violet. Scale bar = 200μm. **F.** Caudal views of cerebella from P21 WT and *Fmnl2^+/418^* mice. SC = superior colliculus; IC = inferior colliculus; H = hemisphere; V = vermis. **G.** Midline sagittal sections of P21 WT and Fmnl2*^418^*^/-^ cerebella stained with hematoxylin and eosin. Lobules are indicated by Roman numerals. Scale bar = 200μm. **H.** Observed proportions of P21 progeny from a cross between *Fmnl2^+/418^* and *Fmnl2^-/-^* mice. Statistical significance was assessed by Chi-square test.

**Fig. S6:**
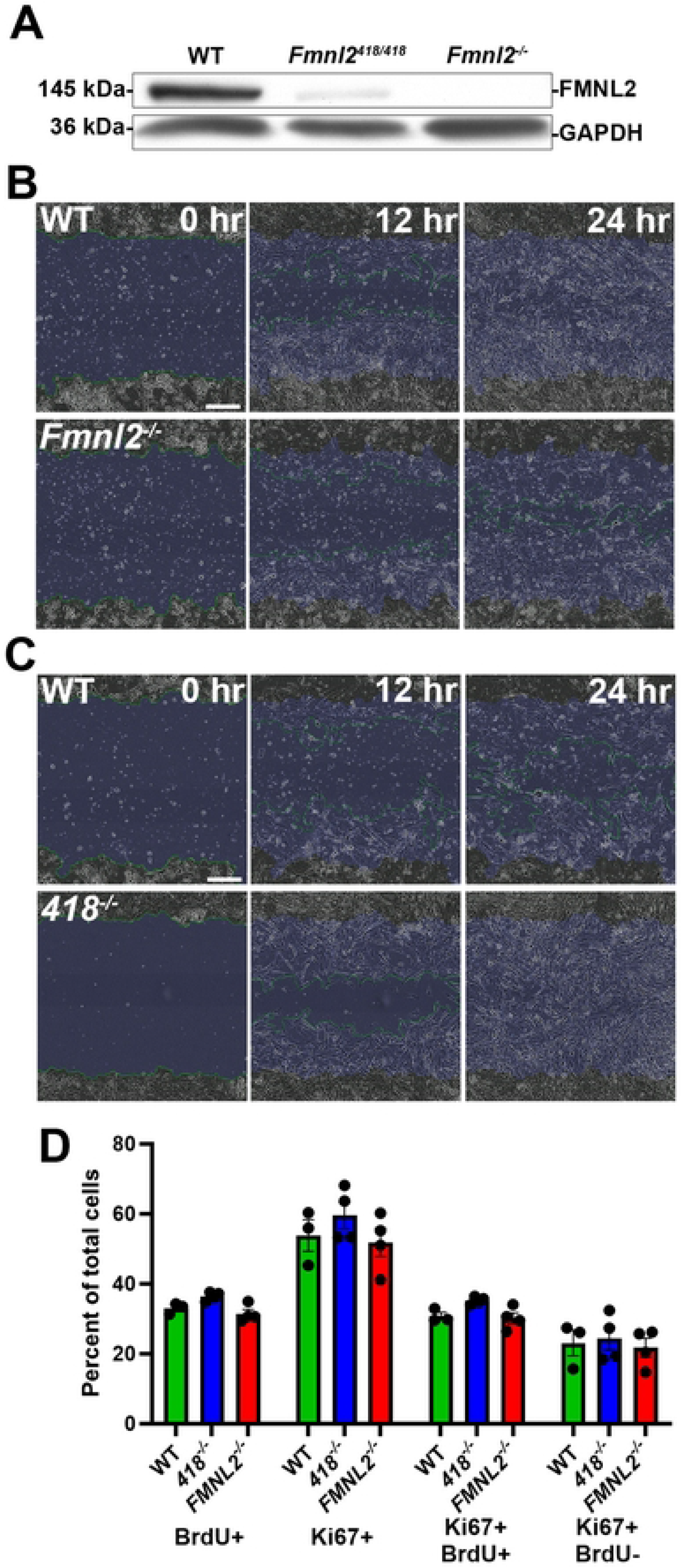
***Fmnl2* deletion or mutation do not alter cell proliferation in mouse embryonic fibroblasts. A.** Representative western blot for FMNL2 in mouse embryonic fibroblasts (MEFs) derived from WT, *Fmnl2^418/418^*, and *Fmnl2*^-/-^ embryos. FMNL2 is markedly reduced in *Fmnl2*^418/418^ MEFs and absent in *Fmnl2*^-/-^ MEFs. GAPDH was used as a loading control. **B, C.** Representative Incucyte images of WT and *Fmnl2*^-/-^ MEF (**B**) and WT and *Fmnl2^418/418^* (**C**) wound healing assays at 0, 12, and 24 hours, corresponding to the quantification in Figure 4H. Scale bars = 100μm. **D.** Quantification of the percentage of WT, *Fmnl2^418/418^*, and *Fmnl2*^-/-^ MEFs positive for BrdU (S-phase), Ki67 (actively cycling), both BrdU and Ki67, or Ki67 alone. Data are means ± SEM. No significant differences in proliferative status were detected across genotypes by two-way ANOVA with Tukey’s multiple comparisons test, confirming that differences in wound closure reflect cell motility rather than proliferation.

**Figure S7:**
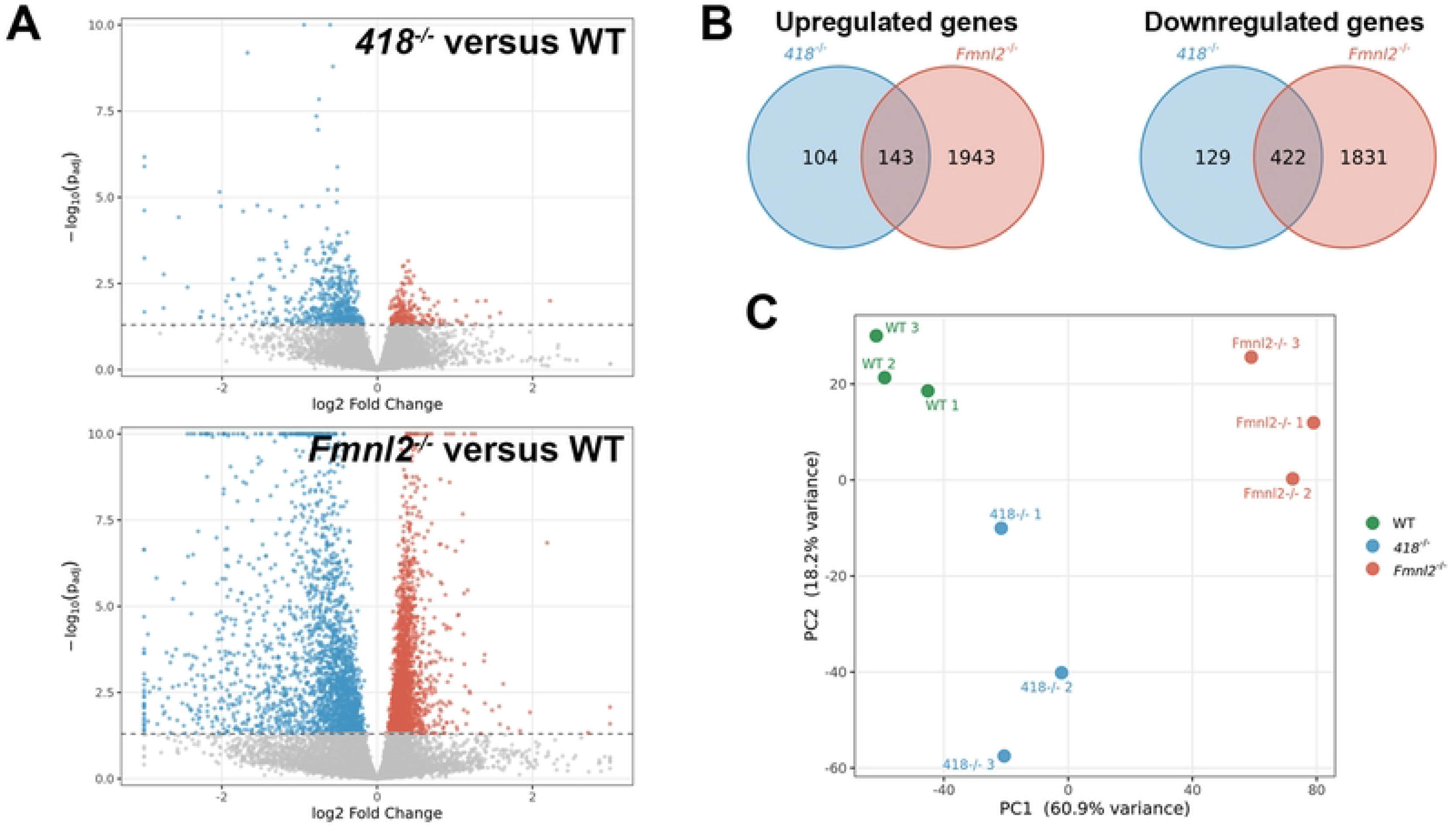
Differential expression analysis and clustering of *Fmnl2^418/418^* and *Fmnl2^-/-^* cerebella. **A.** Volcano plots depicting the spread of gene expression changes between wild-type (WT) and *nmf418^-/-^* or *Fmnl2^-/-^* groups. Differentially expressed genes (p-adj < 0.05) are highlighted in blue if downregulated and red if upregulated in the respective comparisons. **B.** Venn diagrams of the number of upregulated (top) and downregulated (bottom) genes that overlapped between *Fmnl2^418/418^* (*418*^-/-^) and *Fmnl2^-/-^* cerebella when compared to wild-type. **C.** Principal component analysis (PCA) plot of RNA sequencing results depicts clustering of samples by genotype.

